# Identification of essential genes in *Caenorhabditis elegans* through whole genome sequencing of legacy mutant collections

**DOI:** 10.1101/2021.06.16.448744

**Authors:** Erica Li-Leger, Richard Feichtinger, Stephane Flibotte, Heinke Holzkamp, Ralf Schnabel, Donald G. Moerman

## Abstract

It has been estimated that 15-30% of the ∼20,000 genes in *C. elegans* are essential, yet many of these genes remain to be identified or characterized. With the goal of identifying unknown essential genes, we performed whole genome sequencing on complementation pairs from legacy collections of maternal-effect lethal and sterile mutants. This approach uncovered maternal genes required for embryonic development and genes with putative sperm-specific functions. In total, 58 essential genes were identified on chromosomes III, IV, and V, of which 49 genes are represented by novel alleles in this collection. Of these 49 genes, 19 (40 alleles) were selected for further functional characterization. The terminal phenotypes of embryos were examined, revealing defects in cell division, morphogenesis, and osmotic integrity of the eggshell. Mating assays with wild-type males revealed previously unknown male-expressed genes required for fertilization and embryonic development. The result of this study is a catalogue of mutant alleles in essential genes that will serve as a resource to guide further study toward a more complete understanding of this important model organism. As many genes and developmental pathways in *C. elegans* are conserved and essential genes are often linked to human disease, uncovering the function of these genes may also provide insight to further our understanding of human biology.

## INTRODUCTION

Essential genes are those required for the survival or reproduction of an organism, and therefore encode elements that are foundational to life. This class of genes has been widely studied for a number of reasons. Essential genes are often well conserved and can offer insight into the principles that govern common biological processes (Hughes 2002; Jordan *et al*. 2002; Georgi *et al*. 2013). Researching these genes and their functions has important implications in understanding the cellular and developmental processes that form complex organisms, including humans. Additionally, identifying genes that are lethal when mutated opens up new avenues through which drug development approaches can target parasites, pathogens, and cancer cells (for example, Doyle *et al*. 2010; Shi *et al*. 2015; Vyas *et al*. 2015; Zhang *et al*. 2018). Finally, the concept of a minimal gene set that is comprised of all genes necessary for life has been the subject of much investigation and has recently been of particular interest in the field of synthetic biology (reviewed in Ausländer *et al*. 2017).

Studying essential genes in humans is complicated by practical and ethical considerations. Accordingly, model organisms have played a key role in identifying and understanding essential genes, and efforts have been made to identify all essential genes in a few model organisms. Systematic genome-wide studies of gene function in *Saccharomyces cerevisiae* have uncovered more than 1,100 essential genes, many of which have phylogenetically conserved roles in fundamental biological processes such as cell division, protein synthesis and metabolism (Winzeler *et al*. 1999; Giaever *et al*. 2002; Yu *et al*. 2006; Li *et al*. 2011). While an important contribution, this is only a fraction of the all the essential genes in multicellular organisms. In more complex model organisms, identifying all essential genes in the genome has not been so straightforward. The discovery of RNA interference (RNAi; Fire *et al*. 1998) enabled researchers to employ genome-wide reverse genetic screens to examine the phenotypic effects of gene knockdown (Fraser *et al*. 2000; Kamath *et al*. 2003). In general, this has been an effective, high-throughput method for identifying many genes with essential functions (Gönczy *et al*. 2000; Sönnichsen *et al*. 2005). However, there are limitations to using RNAi to screen for all essential genes, including incomplete gene knock down, off-target effects, and RNAi resistance in certain tissue or cell types; thus, many genes of biological importance escape identification in high-throughput RNAi screens. This highlights the motivation to obtain null alleles for every gene in the genome, which has been the goal of several model organism consortia (C. elegans Deletion Mutant Consortium 2012; Bradley *et al*. 2012; Varshney *et al*. 2013), though it has not yet been achieved for any metazoan.

*Caenorhabditis elegans* has been an important model in developmental biology for decades, and the ability to freeze and store populations of *C. elegans* indefinitely allows investigators to share their original mutant strains with others around the world. In the first few decades of *C. elegans* research, dozens of forward genetics screens were used to uncover mutants in hundreds of essential genes (for example, Herman 1978; Meneely and Herman 1979; Rogalski *et al*. 1982; Howell *et al*. 1987; Clark *et al*. 1988; Johnsen and Baillie 1988; Kemphues *et al*. 1988; McKim *et al*. 1988; Howell and Rose 1990; Johnsen and Baillie 1991; McKim *et al*. 1992; Stewart *et al*. 1998; Gönczy *et al*. 1999). These early studies generated what we refer to here as legacy collections. The alleles were often mapped to a region of the genome through deficiency or linkage mapping. However, the process of identifying the molecular nature of the genetic mutations one-by-one using traditional methods was slow and laborious before the genome sequence was complete (The C. elegans Sequencing Consortium 1998) and next-generation sequencing technologies were developed (reviewed in Metzker 2010; Goodwin *et al*. 2016).

As whole genome sequencing (WGS) has become widely adopted, methods for identifying mutant alleles have evolved to take advantage of these technological advances (Sarin *et al*. 2008; Smith *et al*. 2008; Srivatsan *et al*. 2008; Blumenstiel *et al*. 2009; Schneeberger *et al*. 2009; Doitsidou *et al*. 2010; Flibotte *et al*. 2010; Zuryn *et al*. 2010; Smith *et al*. 2016). With WGS becoming increasingly affordable over time, mutant collections can now be mined for data in efficient ways that were not possible two decades ago. Performing WGS on a single mutant genome is often insufficient to identify a causal variant due to the abundance of background mutations in any given strain, particularly one that has been subjected to random mutagenesis (Denver *et al*. 2004; Hillier *et al*. 2008; Sarin *et al*. 2008; Flibotte *et al*. 2010). However, when paired with additional strategies such as deletion or SNP-based mapping or bulk segregant analysis, WGS becomes a valuable tool to expedite gene identification. Furthermore, if multiple independently derived allelic mutants exist, an even simpler approach can be taken. By sequencing two or more mutants within a complementation group and looking for mutations in the same gene, the need for additional mapping or crossing schemes is greatly reduced (Schneeberger *et al*. 2011; Nordström *et al*. 2013).

In the legacy mutant collections described above, where large numbers of mutants are isolated, it is feasible to obtain complementation groups with multiple alleles for many loci. In addition, the abundance of mutants obtained in these large-scale genetic screens suggests that some legacy mutant collections may harbor strains for which the mutations remain unidentified. If such collections are coupled with thorough annotations, they are valuable resources that can be mined with WGS. Indeed, some investigators have recently used such WGS-based approaches to uncover novel essential genes from legacy collections (Jaramillo-Lambert *et al*. 2015; Qin *et al*. 2018). These projects bring us closer to identifying all essential genes in *C. elegans* and also contribute to the ongoing efforts to obtain null mutations in every gene in the genome.

There are currently 3,755 *C. elegans* genes that have been annotated with lethal or sterile phenotypes from RNAi knockdown studies (data from WormBase version WS275). In comparison, the number of genes currently represented by lethal or sterile mutant alleles is 1,885 (data from WormBase version WS275). These numbers should be considered minimums, as the database annotations are not necessarily up to date. The discrepancy in these numbers could be illustrative of the comparatively time-consuming and laborious nature of isolating and identifying mutants. Additionally, some of the genes identified as essential in RNAi screens may belong to paralogous gene families whose redundant functions are masked in single gene knockouts. Although the total number of essential genes in *C. elegans* is unknown, extrapolation from saturation mutagenesis screens has led to estimates that approximately 15-30% of the ∼20,000 genes in this organism are essential (Clark *et al*. 1988; Howell and Rose 1990; Johnsen and Baillie 1997; The C. elegans Deletion Mutant Consortium 2012). This suggests the possibility that there are many essential genes in *C. elegans* that remain unidentified and/or lack representation by a null allele.

In this study, we use WGS to revisit two *C. elegans* legacy mutant collections isolated more than 25 years ago. These collections are a rich resource for essential gene discovery; they comprise 75 complementation groups in which at least two alleles with sterile or maternal-effect lethal phenotypes have been found. With these collections, we sought to identify novel essential genes and to conduct a preliminary characterization of their roles in fertilization and development. Wild-type male rescue assays are used to attribute some mutant phenotypes to sperm-specific genetic defects. In addition, we examine arrested embryos using differential interference contrast (DIC) microscopy and document their terminal phenotypes. This work comprises a catalogue of 123 alleles with mutations in 58 essential genes on chromosomes III, IV, and V. Of these 58 genes, 49 are represented by novel alleles in this collection. We present several genes which are reported here for the first time as essential genes and mutant alleles for genes that have only previously been studied with RNAi knockdown. The aim of this work is to help accelerate research efforts by identifying essential genes and providing an entry point into further investigations of gene function. Advancing our understanding of essential genes is imperative to reaching a more comprehensive knowledge of gene function in *C. elegans* and may provide insight into conserved processes in developmental biology, parasitic nematology, and human disease.

## MATERIALS AND METHODS

### Generation of legacy mutant collections

Mutant strains were isolated in screens for maternal-effect lethal and sterile alleles in the early 1990s by Heinke Holzkamp and Ralf Schnabel (unpublished data), and Richard Feichtinger (Feichtinger 1995). Two balancer strains were used for mutagenesis; GE1532: *unc-32(e189)/qC1 [dpy-19(1259) glp-1(q339)] III; him-3(e1147) IV* and GE1550: *him-9(e1487) II; unc-24(e138)/nT1[let(m435)] IV; dpy-11(e224)/nT1[let(m435)] V.* These parental strains were subjected to ethyl methanesulfonate (EMS) mutagenesis at 20° as described by Brenner (1974), with a mutagen dose of 50-75 mM and duration between 4 and 6 hours. Following mutagenesis, L4 F1 animals were singled on plates at either 15° or 17°. Animals with homozygous markers in the F2 or F3 generation were transferred to 25° and subsequently screened for the production of dead eggs, unfertilized oocytes, or no eggs laid. The two mutant collections analyzed in this study are summarized in Table 1.

**Table 1.** Summary of mutant collections

| Collection | Number of<br>Complementation<br>Groups with $\geq 2$ alleles | Chromosome | Mutant Genotypes |
| --- | --- | --- | --- |
| A | 32 | III | *unc-32(e189) let(t...)/qC1 III; him-3(e1147) IV* |
| B | 25 | IV | *him-9(e1487) II; unc-24(e138) let(t...)/nT1 [let(m435)]*<br>*IV; dpy-11(e224)/nT1 [let(m435)] V* |
|  | 18 | V | *him-9(e1487) II; unc-24(e138)/nT1 [let(m435)] IV; dpy-*<br>*11(e224) let(t...)/nT1 [let(m435)] V* |

### List of strains

The wild-type Bristol N2 derivative PD1074 and strains with the following mutations were used: *him-3(e1147), unc-32(e189), qC1[dpy-19(e1259) glp-1(q339)], him-9(e1487), unc-24(e138), dpy-11(e224, e1180), nT1[let(m435)] (IV;V), nT1[unc(n754)let] (IV;V)*. Strains carrying the following deletions were used for deficiency mapping: *nDf16, nDf40, sDf110, sDf125, tDf5, tDf6, tDf7 (III)*; *eDf19, nDf41, sDf2, sDf21, stDf7 (IV); ctDf1, itDf2, nDf32, sDf28, sDf35 (V)*. All *sDfs* were kindly provided by D. Baillie’s Lab (Simon Fraser University), and some strains were kindly provided by the Caenorhabditis Genetics Center (University of Minnesota). Nematode strains were cultured as previously described by Brenner (1974).

### Outcrossing, mapping and complementation analysis

All mutant strains were outcrossed at least once to minimize background mutations on other chromosomes. Hermaphrodites of the mutant strains were outcrossed with males of GE1532 for Collection A and males of GE1964: *him-9(e1487) II; +/nT1[let(m435)] IV; dpy-11(e1180)/nT1[let(m435)] V* for Collection B. Deficiency mapping was used to localize mutations to a chromosomal region using the deletion strains listed above. A detailed description of the outcrossing and mapping schemes for Collection B can be found in Feichtinger (1995).

Complementation analysis of legacy mutants was performed by crossing 10 males of one mutant strain to 4 hermaphrodites of another strain. The presence of males with homozygous markers indicated successful crossing, and homozygous hermaphrodite progeny were transferred to new plates to determine whether viable offspring were produced and thus complementation occurred. Failure to complement was verified with additional homozygous animals or by repeating the cross. Complementation tests between CRISPR-Cas9 deletion strains and legacy mutants were performed by crossing heterozygous CRISPR-Cas9 deletion (GFP/+) males to heterozygous legacy mutant hermaphrodites. Twenty GFP hermaphrodite F1s were singled on new plates and those segregating viable Dpy and/or Unc progeny indicated complementation between the two alleles.

### DNA extraction

Balanced heterozygous strains were grown on 100 mm nematode growth medium (NGM) agar plates (standard recipe with 3 times concentration of peptone) seeded with OP50 and harvested at starvation. Genomic DNA was extracted using a standard isopropanol precipitation technique previously described (Au *et al*. 2019). DNA quality was assessed with a NanoDrop 2000c Spectrophotometer (Thermo Scientific) and DNA concentration was measured using a Qubit 2.0 Fluorometer and dsDNA Broad Range Assay kit (Life Technologies).

### Whole genome sequencing and analysis pipeline

DNA library preparation and whole genome sequencing were carried out by The Centre for Applied Genomics (The Hospital for Sick Children, Toronto, Canada). Between 20 and 33 *C. elegans* mutant strains were run together on one lane of an Illumina HiSeq X to generate 150-bp paired-end reads.

Sequencing analysis was done using a modified version of a previously designed custom pipeline (Flibotte *et al*. 2010; Thompson *et al*. 2013). Reads were aligned to the *C. elegans* reference genome (WS263; wormbase.org) using the short-read aligner BWA version 0.7.16 (Li and Durbin 2009). Single nucleotide variants (SNVs) and small insertions or deletions (indels) were called using SAMtools toolbox version 1.6 (Li *et al*. 2009). To eliminate unreliable calls, variants at genomic locations for which the canonical N2 strain has historically had low read depth or poor quality (Thompson *et al*. 2013) were removed as potential candidates. The variant calls were annotated with a custom Perl script and labeled heterozygous if represented by 20-80% of the reads at that location. The remaining candidates were then subjected to a series of custom filters. Any variants that appeared in more than three strains from the same collection were removed. The remaining list was filtered to only include heterozygous mutations affecting coding exons (indels, missense and nonsense mutations) and splice sites (defined as the first two and last two base pairs in an intron). Finally, the list of candidate mutations was trimmed to include only mutations on the chromosome to which the mutation had originally been mapped.

For each pair of strains belonging to a complementation group, the final list of candidate mutations was compared and the gene or genes in common were identified. In cases where there was only one gene in common on both lists, this gene was designated the candidate essential gene. For complementation groups with multiple candidate genes in common, additional information such as the nature of the mutations and existing knowledge about the genes was used to select a single candidate gene, when possible. When there was no gene candidate in common within a pair of strains, the list of variants was reanalyzed to look for larger deletions and rearrangements. If available, two additional alleles were sequenced to help identify the gene.

### Validation of gene identities

To validate the candidate gene identities derived from whole genome sequencing analysis, the genomic position of each candidate gene was corroborated with the legacy data from deficiency mapping experiments. Approximate boundaries for the deletions were estimated from the map coordinates of genes known to lie internal or external to the deletions according to data from WormBase (WS275).

For further validation of select gene candidates, deletion mutants were generated in an N2 wild-type background using a CRISPR-Cas9 genome editing strategy previously described (Norris *et al*. 2015; Au *et al*. 2019). Two guide RNAs were used to excise the gene of interest and replace it with a selection cassette expressing G418 drug resistance and pharyngeal GFP (*loxP* + P*myo-2*::*GFP*::*unc-54* 3′UTR + P*rps-27*::*neoR*::*unc-54*3′UTR *+ loxP* vector, provided by Dr. John Calarco, University of Toronto, Canada). Guide RNAs were designed using the *C. elegans* Guide Selection Tool (genome.sfu.ca/crispr) and synthesized by Integrated DNA Technologies (IDT). Repair templates were generated by assembling homology arms (450-bp gBlocks synthesized by IDT) and the selection cassette using the NEBuilder Hifi DNA Assembly Kit (New England Biolabs).

Cas9 protein (generously gifted from Dr. Geraldine Seydoux) was assembled into a ribonucleoprotein (RNP) complex with the guide RNAs and tracrRNA (IDT) following the manufacturer’s recommendations. PD1074 animals were injected using standard microinjection techniques (Mello *et al*. 1991; Kadandale *et al*. 2009) with an injection mix consisting of: 50 ng/µl repair template, 0.5 µM RNP complex, 5 ng/µl pCFJ104 (P*myo-3*::mCherry), and 2.5 ng/µl pCFJ90 (P*myo-2*::mCherry). Injected animals were screened according to the protocol described in Norris *et al*. (2015) and genomic edits were validated using the PCR protocol described in Au *et al*. (2019). Complementation tests between CRISPR-Cas9 alleles and legacy mutant alleles were performed to verify gene identities, as described above.

### Analysis of orthologs, gene ontology, and expression patterns

Previously reported phenotypes from RNAi experiments or mutant alleles were retrieved from WormBase (WS275) and GExplore (genome.sfu.ca/gexplore; Hutter *et al*. 2009; Hutter and Suh 2016). Life stage-specific gene expression data from the modENCODE project (Hillier *et al*. 2009; Gerstein *et al*. 2010, 2014; Boeck *et al*. 2016) were also accessed through GExplore. Visual inspection of these data revealed genes with maternal expression patterns (high levels of expression in the early embryo and hermaphrodite gonad) as well as those predominantly expressed in males.

Human orthologs of *C. elegans* genes were determined using Ortholist 2 (ortholist.shaye-lab.org; Kim *et al*. 2018). For maximum sensitivity, the minimum number of programs predicting a given ortholog was set to one. NCBI BLASTp (blast.ncbi.nlm.nih.gov; Altschul *et al*. 1990) was used to examine distributions of homologs across species and potential nematode-specificity in genes with no human orthologs. Protein sequences from the longest transcript of each gene were used to query the non-redundant protein sequences (nr) database, with default parameters and a maximum of 1,000 target sequences. The results were filtered with an E-value threshold of 10^-5^.

Gene Ontology (GO) term analysis was performed using PANTHER version 16.0 (Thomas *et al*. 2003). The list of 58 candidate genes was used for an overrepresentation test, with the set of all *C. elegans* genes as a background list. Overrepresentation was analyzed with a Fisher’s Exact test and p-values were adjusted with the Bonferroni multiple testing correction.

### Temperature sensitivity and mating assays

To assay temperature sensitivity, heterozygous strains were propagated at 15° and homozygous L4 animals were isolated on 60 mm NGM plates (2 x 6/plate or 3 x 3/plate). After one week at 15°, plates were screened for the presence of viable homozygous progeny. If present, L4 homozygotes were transferred to new plates at 25° and screened after three days to confirm lethality or sterility.

Mating assays were carried out using PD1074 males and mutant hermaphrodites. Three L4-stage homozygous mutant hermaphrodites were isolated and crossed with ten PD1074 males on each of three 60 mm NGM plates. Control plates consisted of three L4 hermaphrodite mutants without males. Mating assays were carried out at 25°C and observations were taken after three days, noting the absence or presence of viable cross progeny.

### Microscopy

The terminal phenotypes of dead eggs from maternal-effect lethal mutants were observed using DIC microscopy. Young adult homozygous mutants were dissected to release their eggs in either M9 buffer with Triton X-100 (0.5%; M9+TX) or distilled water and embryos were left to develop at 25°C overnight (∼16 hours). Embryos were mounted on 2% agarose pads and visualized using a Zeiss Axioplan 2 equipped with DIC optics. Images of representative embryos were captured using a Zeiss Axiocam 105 Color camera and ZEN 2.6 imaging software (Carl Zeiss Microscopy). For embryos incubated in distilled water, an osmotic integrity defective (OID) phenotype was noted for embryos that burst or swelled and filled the eggshell, as described by Sönnichsen *et al*. (2005).

### Data availability

The raw sequence data from this study have been deposited in the NCBI Sequence Read Archive (SRA; ncbi.nlm.nih.gov/sra) under accession number PRJNA628853. Supplemental material is available at Figshare. File S1 contains sequences and associated information for CRISPR-Cas9 deletion alleles. File S2 contains life stage-specific expression patterns for the Genes of Interest. File S3 contains documentation of the terminal phenotypes for maternal-effect lethal embryos.

## RESULTS

### Identification of 58 essential genes

Whole genome sequencing was performed on a total of 157 strains, with depth of coverage ranging between 21x and 65x (average = 38x). A minimum of two alleles for each of 75 complementation groups were sequenced and a total of 58 essential genes were identified (Table 2). Literature searches revealed that 43 of these genes have been annotated with lethal or sterile phenotypes from either mutant alleles or RNAi studies. Furthermore, 17 of the 157 alleles had been previously sequenced (Vatcher *et al*. 1998; Gönczy *et al*. 2001; Kaitna *et al*. 2002; Brauchle *et al*. 2003; Cockell *et al*. 2004; Delattre *et al*. 2004; Sonneville *et al*. 2004; Bischoff and Schnabel 2006; Nieto *et al*. 2010), and therefore served as a blind test set to validate our analysis approach. Eight of the nine genes represented in this blind test set were correctly identified by our pipeline, whereas one gene escaped identification. This was due to an intronic mutation that did not pass our filtering criteria but was found upon manual inspection of the sequencing data. While the list of 58 genes includes many known essential genes, among the known genes are alleles that are novel genetic variants. Nineteen genes from this collection which were not previously studied or were not represented by lethal or sterile mutants were designated Genes of Interest (GOI; Table 3). These 19 GOI, represented by 40 alleles, were further characterized as part of this study. They include 14 genes (28 alleles) with a maternal-effect lethal phenotype and 5 genes (12 alleles) with a sterile phenotype.

**Table 2.**
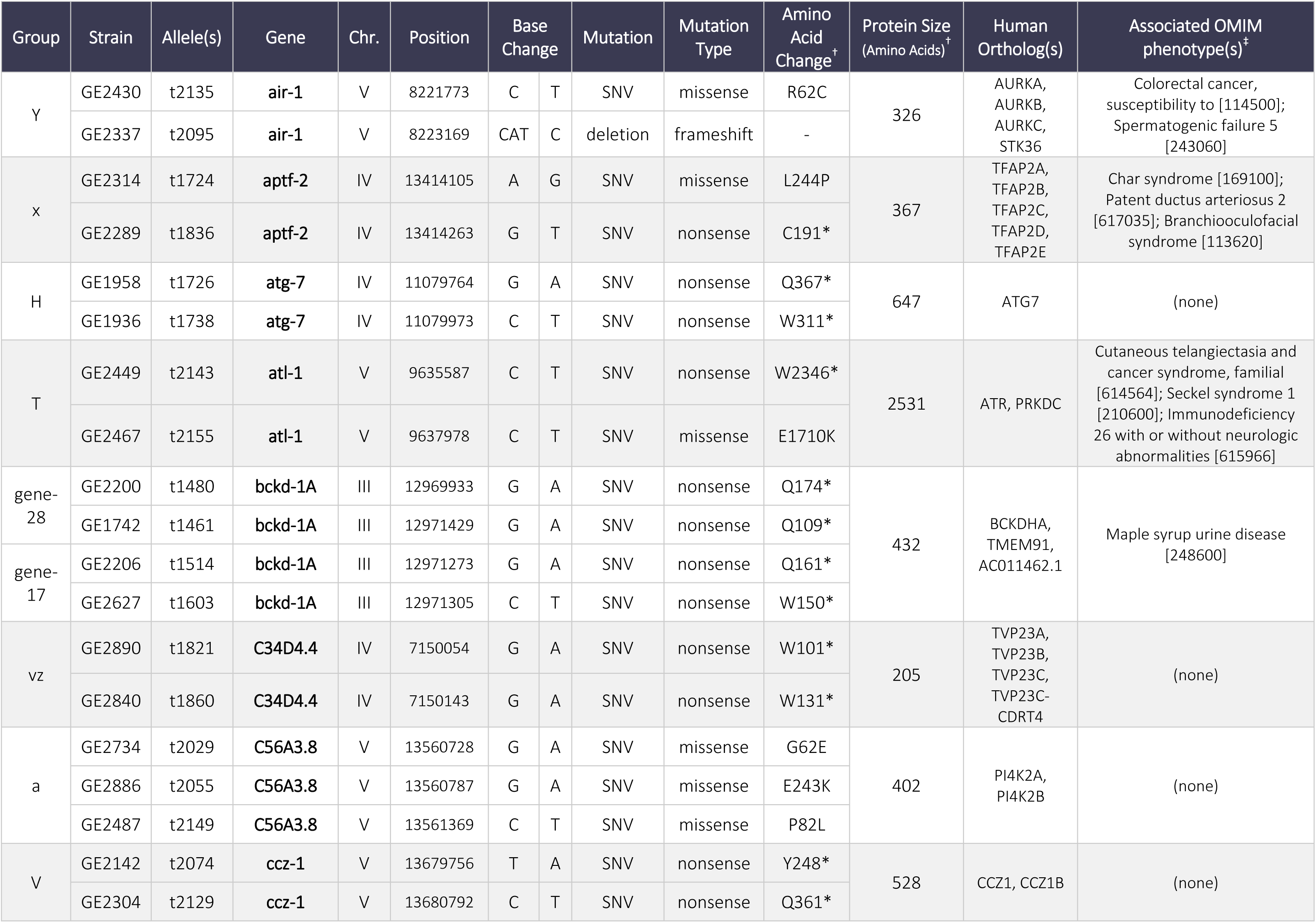
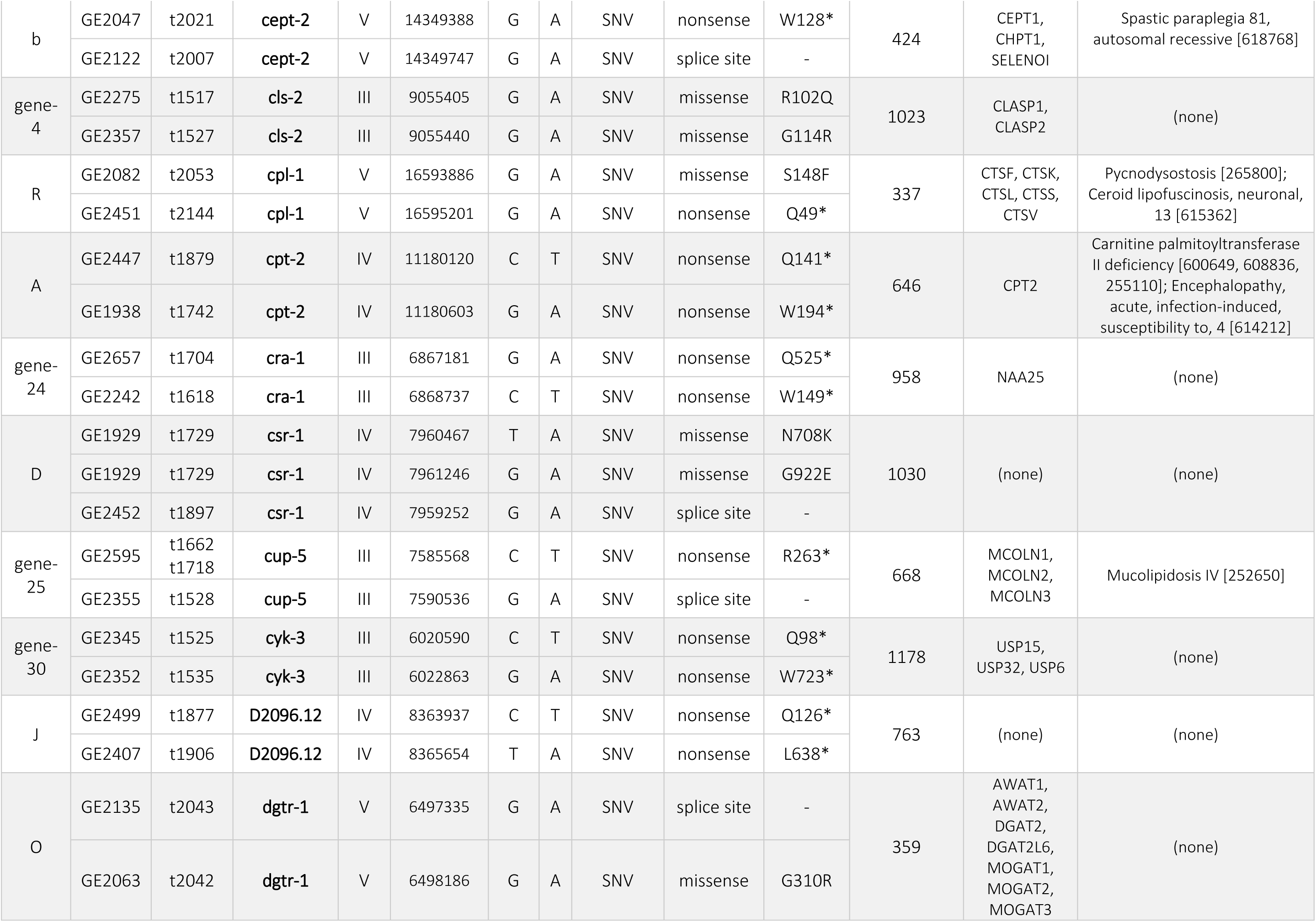
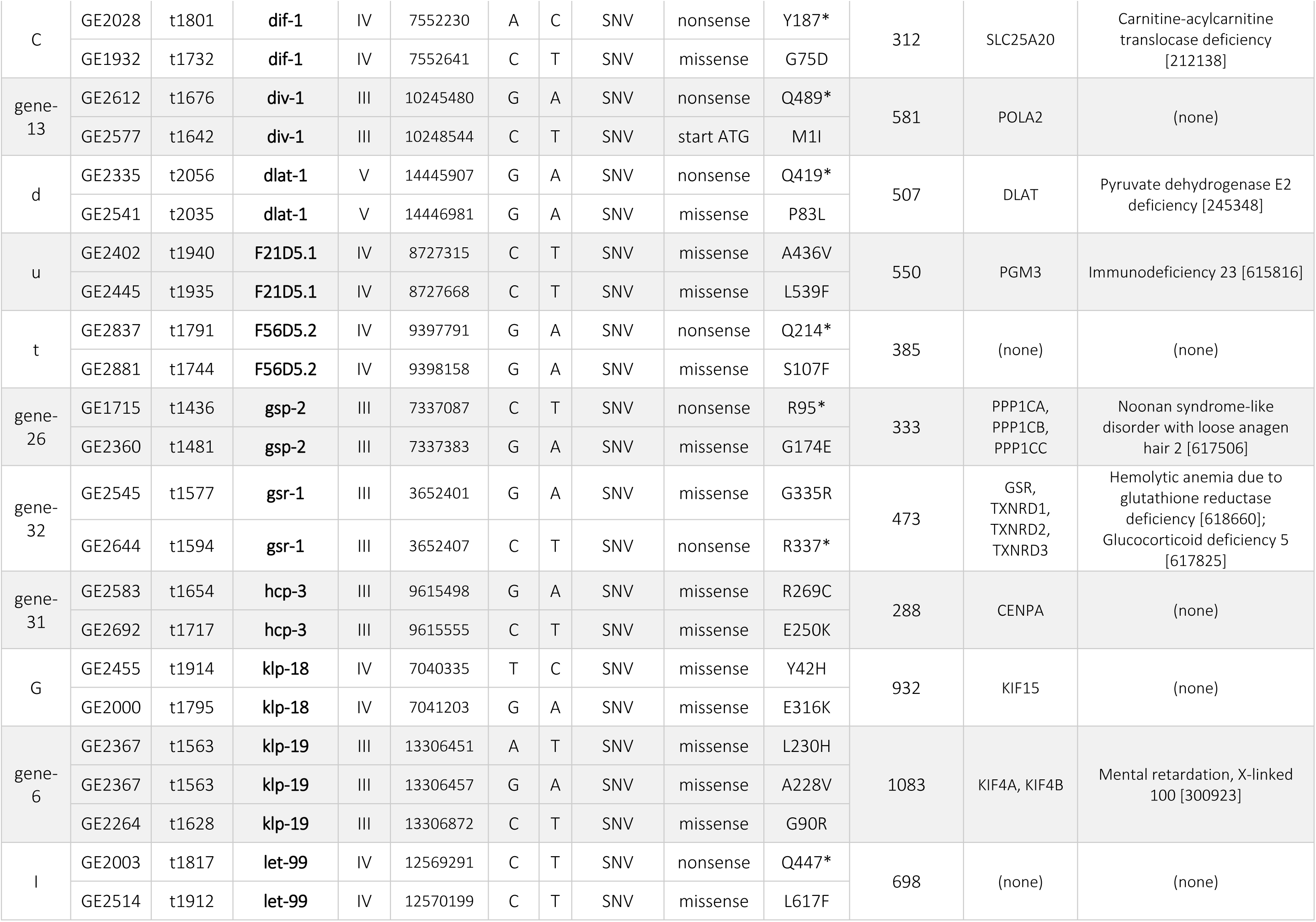
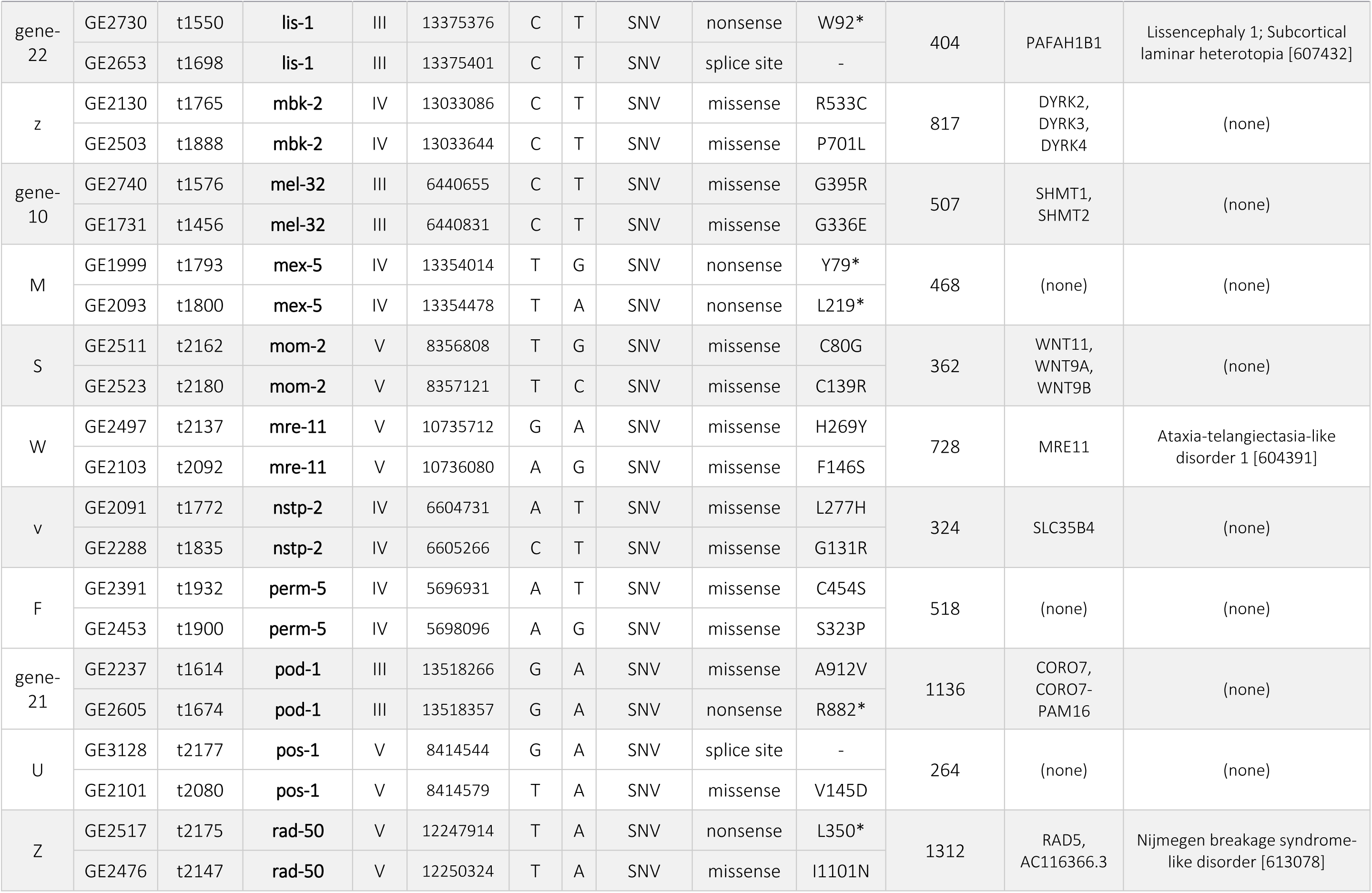
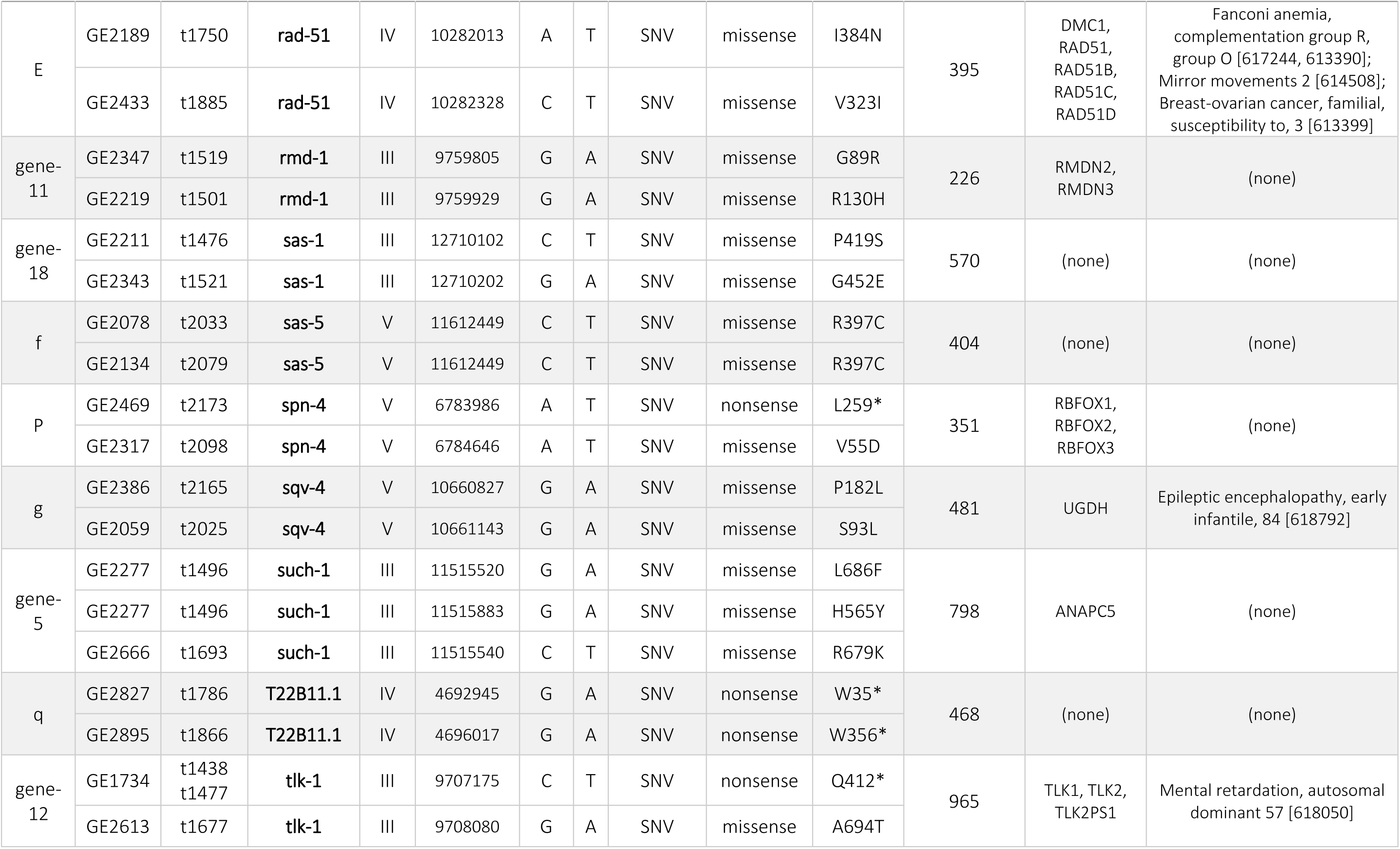
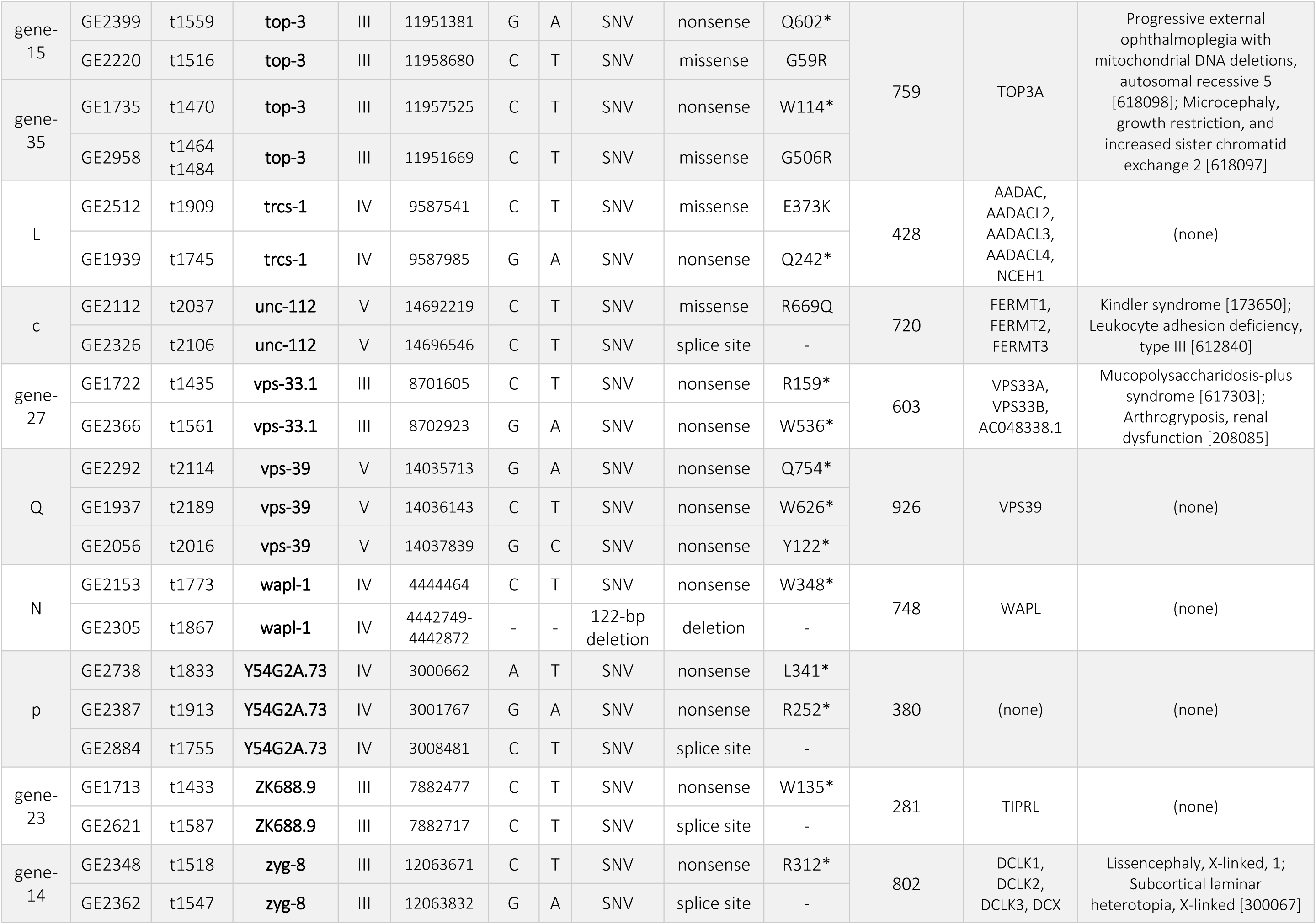
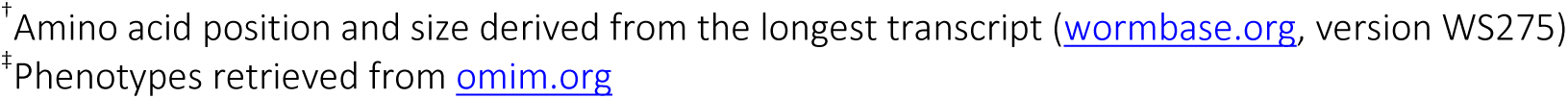
List of 58 essential genes with associated maternal-effect lethal or sterile alleles

| Group | Strain | Allele(s) | Gene | Chr. | Position | Base Change |  | Mutation | Mutation Type | Amino Acid Change<sup>†</sup> | Protein Size (Amino Acids)<sup>†</sup> | Human Ortholog(s) | Associated OMIM phenotype(s)<sup>‡</sup> |
| --- | --- | --- | --- | --- | --- | --- | --- | --- | --- | --- | --- | --- | --- |
| Y | GE2430 | t2135 | **air-1** | V | 8221773 | C | T | SNV | missense | R62C | 326 | AURKA, AURKB, AURKC, STK36 | Colorectal cancer, susceptibility to [114500]; Spermatogenic failure 5 [243060] |
|  | GE2337 | t2095 | **air-1** | V | 8223169 | CAT | C | deletion | frameshift | - |  |  |  |
| x | GE2314 | t1724 | **aptf-2** | IV | 13414105 | A | G | SNV | missense | L244P | 367 | TFAP2A, TFAP2B, TFAP2C, TFAP2D, TFAP2E | Char syndrome [169100]; Patent ductus arteriosus 2 [617035]; Branchiooculofacial syndrome [113620] |
|  | GE2289 | t1836 | **aptf-2** | IV | 13414263 | G | T | SNV | nonsense | C191* |  |  |  |
| H | GE1958 | t1726 | **atg-7** | IV | 11079764 | G | A | SNV | nonsense | Q367* | 647 | ATG7 | (none) |
|  | GE1936 | t1738 | **atg-7** | IV | 11079973 | C | T | SNV | nonsense | W311* |  |  |  |
| T | GE2449 | t2143 | **atl-1** | V | 9635587 | C | T | SNV | nonsense | W2346* | 2531 | ATR, PRKDC | Cutaneous telangiectasia and cancer syndrome, familial [614564]; Seckel syndrome 1 [210600]; Immunodeficiency 26 with or without neurologic abnormalities [615966] |
|  | GE2467 | t2155 | **atl-1** | V | 9637978 | C | T | SNV | missense | E1710K |  |  |  |
| gene-28 | GE2200 | t1480 | **bckd-1A** | III | 12969933 | G | A | SNV | nonsense | Q174* | 432 | BCKDHA, TMEM91, AC011462.1 | Maple syrup urine disease [248600] |
|  | GE1742 | t1461 | **bckd-1A** | III | 12971429 | G | A | SNV | nonsense | Q109* |  |  |  |
| gene-17 | GE2206 | t1514 | **bckd-1A** | III | 12971273 | G | A | SNV | nonsense | Q161* |  |  |  |
|  | GE2627 | t1603 | **bckd-1A** | III | 12971305 | C | T | SNV | nonsense | W150* |  |  |  |
| vz | GE2890 | t1821 | **C34D4.4** | IV | 7150054 | G | A | SNV | nonsense | W101* | 205 | TVP23A, TVP23B, TVP23C, TVP23C-CDRT4 | (none) |
|  | GE2840 | t1860 | **C34D4.4** | IV | 7150143 | G | A | SNV | nonsense | W131* |  |  |  |
| a | GE2734 | t2029 | **C56A3.8** | V | 13560728 | G | A | SNV | missense | G62E | 402 | PI4K2A, PI4K2B | (none) |
|  | GE2886 | t2055 | **C56A3.8** | V | 13560787 | G | A | SNV | missense | E243K |  |  |  |
|  | GE2487 | t2149 | **C56A3.8** | V | 13561369 | C | T | SNV | missense | P82L |  |  |  |
| V | GE2142 | t2074 | **ccz-1** | V | 13679756 | T | A | SNV | nonsense | Y248* | 528 | CCZ1, CCZ1B | (none) |
|  | GE2304 | t2129 | **ccz-1** | V | 13680792 | C | T | SNV | nonsense | Q361* |  |  |  |
| b | GE2047 | t2021 | **cept-2** | V | 14349388 | G | A | SNV | nonsense | W128* | 424 | CEPT1, CHPT1, SELENOI | Spastic paraplegia 81, autosomal recessive [618768] |
|  | GE2122 | t2007 | **cept-2** | V | 14349747 | G | A | SNV | splice site | - |  |  |  |
| gene-4 | GE2275 | t1517 | **cls-2** | III | 9055405 | G | A | SNV | missense | R102Q | 1023 | CLASP1, CLASP2 | (none) |
|  | GE2357 | t1527 | **cls-2** | III | 9055440 | G | A | SNV | missense | G114R |  |  |  |
| R | GE2082 | t2053 | **cpl-1** | V | 16593886 | G | A | SNV | missense | S148F | 337 | CTSF, CTSK, CTSL, CTSS, CTSV | Pycnodysostosis [265800]; Ceroid lipofuscinosis, neuronal, 13 [615362] |
|  | GE2451 | t2144 | **cpl-1** | V | 16595201 | G | A | SNV | nonsense | Q49* |  |  |  |
| A | GE2447 | t1879 | **cpt-2** | IV | 11180120 | C | T | SNV | nonsense | Q141* | 646 | CPT2 | Carnitine palmitoyltransferase II deficiency [600649, 608836, 255110]; Encephalopathy, acute, infection-induced, susceptibility to, 4 [614212] |
|  | GE1938 | t1742 | **cpt-2** | IV | 11180603 | G | A | SNV | nonsense | W194* |  |  |  |
| gene-24 | GE2657 | t1704 | **cra-1** | III | 6867181 | G | A | SNV | nonsense | Q525* | 958 | NAA25 | (none) |
|  | GE2242 | t1618 | **cra-1** | III | 6868737 | C | T | SNV | nonsense | W149* |  |  |  |
| D | GE1929 | t1729 | **csr-1** | IV | 7960467 | T | A | SNV | missense | N708K | 1030 | (none) | (none) |
|  | GE1929 | t1729 | **csr-1** | IV | 7961246 | G | A | SNV | missense | G922E |  |  |  |
|  | GE2452 | t1897 | **csr-1** | IV | 7959252 | G | A | SNV | splice site | - |  |  |  |
| gene-25 | GE2595 | t1662<br>t1718 | **cup-5** | III | 7585568 | C | T | SNV | nonsense | R263* | 668 | MCOLN1, MCOLN2, MCOLN3 | Mucopolipidosis IV [252650] |
|  | GE2355 | t1528 | **cup-5** | III | 7590536 | G | A | SNV | splice site | - |  |  |  |
| gene-30 | GE2345 | t1525 | **cyk-3** | III | 6020590 | C | T | SNV | nonsense | Q98* | 1178 | USP15, USP32, USP6 | (none) |
|  | GE2352 | t1535 | **cyk-3** | III | 6022863 | G | A | SNV | nonsense | W723* |  |  |  |
| J | GE2499 | t1877 | **D2096.12** | IV | 8363937 | C | T | SNV | nonsense | Q126* | 763 | (none) | (none) |
|  | GE2407 | t1906 | **D2096.12** | IV | 8365654 | T | A | SNV | nonsense | L638* |  |  |  |
| O | GE2135 | t2043 | **dgtr-1** | V | 6497335 | G | A | SNV | splice site | - | 359 | AWAT1, AWAT2, DGAT2, DGAT2L6, MOGAT1, MOGAT2, MOGAT3 | (none) |
|  | GE2063 | t2042 | **dgtr-1** | V | 6498186 | G | A | SNV | missense | G310R |  |  |  |
| C | GE2028 | t1801 | **dif-1** | IV | 7552230 | A | C | SNV | nonsense | Y187* | 312 | SLC25A20 | Carnitine-acylcarnitine translocase deficiency [212138] |
|  | GE1932 | t1732 | **dif-1** | IV | 7552641 | C | T | SNV | missense | G75D |  |  |  |
| gene-13 | GE2612 | t1676 | **div-1** | III | 10245480 | G | A | SNV | nonsense | Q489* | 581 | POLA2 | (none) |
|  | GE2577 | t1642 | **div-1** | III | 10248544 | C | T | SNV | start ATG | M1I |  |  |  |
| d | GE2335 | t2056 | **dlat-1** | V | 14445907 | G | A | SNV | nonsense | Q419* | 507 | DLAT | Pyruvate dehydrogenase E2 deficiency [245348] |
|  | GE2541 | t2035 | **dlat-1** | V | 14446981 | G | A | SNV | missense | P83L |  |  |  |
| u | GE2402 | t1940 | **F21D5.1** | IV | 8727315 | C | T | SNV | missense | A436V | 550 | PGM3 | Immunodeficiency 23 [615816] |
|  | GE2445 | t1935 | **F21D5.1** | IV | 8727668 | C | T | SNV | missense | L539F |  |  |  |
| t | GE2837 | t1791 | **F56D5.2** | IV | 9397791 | G | A | SNV | nonsense | Q214* | 385 | (none) | (none) |
|  | GE2881 | t1744 | **F56D5.2** | IV | 9398158 | G | A | SNV | missense | S107F |  |  |  |
| gene-26 | GE1715 | t1436 | **gsp-2** | III | 7337087 | C | T | SNV | nonsense | R95* | 333 | PPP1CA, PPP1CB, PPP1CC | Noonan syndrome-like disorder with loose anagen hair 2 [617506] |
|  | GE2360 | t1481 | **gsp-2** | III | 7337383 | G | A | SNV | missense | G174E |  |  |  |
| gene-32 | GE2545 | t1577 | **gsr-1** | III | 3652401 | G | A | SNV | missense | G335R | 473 | GSR, TXNRD1, TXNRD2, TXNRD3 | Hemolytic anemia due to glutathione reductase deficiency [618660]; Glucocorticoid deficiency 5 [617825] |
|  | GE2644 | t1594 | **gsr-1** | III | 3652407 | C | T | SNV | nonsense | R337* |  |  |  |
| gene-31 | GE2583 | t1654 | **hcp-3** | III | 9615498 | G | A | SNV | missense | R269C | 288 | CENPA | (none) |
|  | GE2692 | t1717 | **hcp-3** | III | 9615555 | C | T | SNV | missense | E250K |  |  |  |
| G | GE2455 | t1914 | **klp-18** | IV | 7040335 | T | C | SNV | missense | Y42H | 932 | KIF15 | (none) |
|  | GE2000 | t1795 | **klp-18** | IV | 7041203 | G | A | SNV | missense | E316K |  |  |  |
| gene-6 | GE2367 | t1563 | **klp-19** | III | 13306451 | A | T | SNV | missense | L230H | 1083 | KIF4A, KIF4B | Mental retardation, X-linked 100 [300923] |
|  | GE2367 | t1563 | **klp-19** | III | 13306457 | G | A | SNV | missense | A228V |  |  |  |
|  | GE2264 | t1628 | **klp-19** | III | 13306872 | C | T | SNV | missense | G90R |  |  |  |
| I | GE2003 | t1817 | **let-99** | IV | 12569291 | C | T | SNV | nonsense | Q447* | 698 | (none) | (none) |
|  | GE2514 | t1912 | **let-99** | IV | 12570199 | C | T | SNV | missense | L617F |  |  |  |

| Group | Strain | Allele(s) | Gene | Chr. | Position | Base Change |  | Mutation | Mutation Type | Amino Acid Change<sup>†</sup> | Protein Size (Amino Acids)<sup>†</sup> | Human Ortholog(s) | Associated OMIM phenotype(s)<sup>†</sup> |
| --- | --- | --- | --- | --- | --- | --- | --- | --- | --- | --- | --- | --- | --- |
| gene-22 | GE2730 | t1550 | **lis-1** | III | 13375376 | C | T | SNV | nonsense | W92* | 404 | PAFAH1B1 | Lissencephaly 1; Subcortical laminar heterotopia [607432] |
|  | GE2653 | t1698 | **lis-1** | III | 13375401 | C | T | SNV | splice site | - |  |  |  |
| z | GE2130 | t1765 | **mbk-2** | IV | 13033086 | C | T | SNV | missense | R533C | 817 | DYRK2, DYRK3, DYRK4 | (none) |
|  | GE2503 | t1888 | **mbk-2** | IV | 13033644 | C | T | SNV | missense | P701L |  |  |  |
| gene-10 | GE2740 | t1576 | **mel-32** | III | 6440655 | C | T | SNV | missense | G395R | 507 | SHMT1, SHMT2 | (none) |
|  | GE1731 | t1456 | **mel-32** | III | 6440831 | C | T | SNV | missense | G336E |  |  |  |
| M | GE1999 | t1793 | **mex-5** | IV | 13354014 | T | G | SNV | nonsense | Y79* | 468 | (none) | (none) |
|  | GE2093 | t1800 | **mex-5** | IV | 13354478 | T | A | SNV | nonsense | L219* |  |  |  |
| S | GE2511 | t2162 | **mom-2** | V | 8356808 | T | G | SNV | missense | C80G | 362 | WNT11, WNT9A, WNT9B | (none) |
|  | GE2523 | t2180 | **mom-2** | V | 8357121 | T | C | SNV | missense | C139R |  |  |  |
| W | GE2497 | t2137 | **mre-11** | V | 10735712 | G | A | SNV | missense | H269Y | 728 | MRE11 | Ataxia-telangiectasia-like disorder 1 [604391] |
|  | GE2103 | t2092 | **mre-11** | V | 10736080 | A | G | SNV | missense | F146S |  |  |  |
| v | GE2091 | t1772 | **nstp-2** | IV | 6604731 | A | T | SNV | missense | L277H | 324 | SLC35B4 | (none) |
|  | GE2288 | t1835 | **nstp-2** | IV | 6605266 | C | T | SNV | missense | G131R |  |  |  |
| F | GE2391 | t1932 | **perm-5** | IV | 5696931 | A | T | SNV | missense | C454S | 518 | (none) | (none) |
|  | GE2453 | t1900 | **perm-5** | IV | 5698096 | A | G | SNV | missense | S323P |  |  |  |
| gene-21 | GE2237 | t1614 | **pod-1** | III | 13518266 | G | A | SNV | missense | A912V | 1136 | CORO7, CORO7-PAM16 | (none) |
|  | GE2605 | t1674 | **pod-1** | III | 13518357 | G | A | SNV | nonsense | R882* |  |  |  |
| U | GE3128 | t2177 | **pos-1** | V | 8414544 | G | A | SNV | splice site | - | 264 | (none) | (none) |
|  | GE2101 | t2080 | **pos-1** | V | 8414579 | T | A | SNV | missense | V145D |  |  |  |
| Z | GE2517 | t2175 | **rad-50** | V | 12247914 | T | A | SNV | nonsense | L350* | 1312 | RAD5, AC116366.3 | Nijmegen breakage syndrome-like disorder [613078] |
|  | GE2476 | t2147 | **rad-50** | V | 12250324 | T | A | SNV | missense | I1101N |  |  |  |
| E | GE2189 | t1750 | **rad-51** | IV | 10282013 | A | T | SNV | missense | I384N | 395 | DMC1, RAD51, RAD51B, RAD51C, RAD51D | Fanconi anemia, complementation group R, group O [617244, 613390]; Mirror movements 2 [614508]; Breast-ovarian cancer, familial, susceptibility to, 3 [613399] |
|  | GE2433 | t1885 | **rad-51** | IV | 10282328 | C | T | SNV | missense | V323I |  |  |  |
| gene-11 | GE2347 | t1519 | **rmd-1** | III | 9759805 | G | A | SNV | missense | G89R | 226 | RMDN2, RMDN3 | (none) |
|  | GE2219 | t1501 | **rmd-1** | III | 9759929 | G | A | SNV | missense | R130H |  |  |  |
| gene-18 | GE2211 | t1476 | **sas-1** | III | 12710102 | C | T | SNV | missense | P419S | 570 | (none) | (none) |
|  | GE2343 | t1521 | **sas-1** | III | 12710202 | G | A | SNV | missense | G452E |  |  |  |
| f | GE2078 | t2033 | **sas-5** | V | 11612449 | C | T | SNV | missense | R397C | 404 | (none) | (none) |
|  | GE2134 | t2079 | **sas-5** | V | 11612449 | C | T | SNV | missense | R397C |  |  |  |
| P | GE2469 | t2173 | **spn-4** | V | 6783986 | A | T | SNV | nonsense | L259* | 351 | RBFOX1, RBFOX2, RBFOX3 | (none) |
|  | GE2317 | t2098 | **spn-4** | V | 6784646 | A | T | SNV | missense | V55D |  |  |  |
| g | GE2386 | t2165 | **sqv-4** | V | 10660827 | G | A | SNV | missense | P182L | 481 | UGDH | Epileptic encephalopathy, early infantile, 84 [618792] |
|  | GE2059 | t2025 | **sqv-4** | V | 10661143 | G | A | SNV | missense | S93L |  |  |  |
| gene-5 | GE2277 | t1496 | **such-1** | III | 11515520 | G | A | SNV | missense | L686F | 798 | ANAPC5 | (none) |
|  | GE2277 | t1496 | **such-1** | III | 11515883 | G | A | SNV | missense | H565Y |  |  |  |
|  | GE2666 | t1693 | **such-1** | III | 11515540 | C | T | SNV | missense | R679K |  |  |  |
| q | GE2827 | t1786 | **T22B11.1** | IV | 4692945 | G | A | SNV | nonsense | W35* | 468 | (none) | (none) |
|  | GE2895 | t1866 | **T22B11.1** | IV | 4696017 | G | A | SNV | nonsense | W356* |  |  |  |
| gene-12 | GE1734 | t1438<br>t1477 | **tlk-1** | III | 9707175 | C | T | SNV | nonsense | Q412* | 965 | TLK1, TLK2, TLK2PS1 | Mental retardation, autosomal dominant 57 [618050] |
|  | GE2613 | t1677 | **tlk-1** | III | 9708080 | G | A | SNV | missense | A694T |  |  |  |

| Group | Strain | Allele(s) | Gene | Chr. | Position | Base Change |  | Mutation | Mutation Type | Amino Acid Change<sup>†</sup> | Protein Size (Amino Acids)<sup>‡</sup> | Human Ortholog(s) | Associated OMIM phenotype(s)<sup>‡</sup> |
| --- | --- | --- | --- | --- | --- | --- | --- | --- | --- | --- | --- | --- | --- |
| gene-15 | GE2399 | t1559 | **top-3** | III | 11951381 | G | A | SNV | nonsense | Q602* | 759 | TOP3A | Progressive external ophthalmoplegia with mitochondrial DNA deletions, autosomal recessive 5 [618098]; Microcephaly, growth restriction, and increased sister chromatid exchange 2 [618097] |
|  | GE2220 | t1516 | **top-3** | III | 11958680 | C | T | SNV | missense | G59R |  |  |  |
| gene-35 | GE1735 | t1470 | **top-3** | III | 11957525 | C | T | SNV | nonsense | W114* |  |  |  |
|  | GE2958 | t1464<br>t1484 | **top-3** | III | 11951669 | C | T | SNV | missense | G506R |  |  |  |
| L | GE2512 | t1909 | **trcs-1** | IV | 9587541 | C | T | SNV | missense | E373K | 428 | AADAC, AADACL2, AADACL3, AADACL4, NCEH1 | (none) |
|  | GE1939 | t1745 | **trcs-1** | IV | 9587985 | G | A | SNV | nonsense | Q242* |  |  |  |
| c | GE2112 | t2037 | **unc-112** | V | 14692219 | C | T | SNV | missense | R669Q | 720 | FERMT1, FERMT2, FERMT3 | Kindler syndrome [173650]; Leukocyte adhesion deficiency, type III [612840] |
|  | GE2326 | t2106 | **unc-112** | V | 14696546 | C | T | SNV | splice site | - |  |  |  |
| gene-27 | GE1722 | t1435 | **vps-33.1** | III | 8701605 | C | T | SNV | nonsense | R159* | 603 | VPS33A, VPS33B, AC048338.1 | Mucopolysaccharidosis-plus syndrome [617303]; Arthrogryposis, renal dysfunction [208085] |
|  | GE2366 | t1561 | **vps-33.1** | III | 8702923 | G | A | SNV | nonsense | W536* |  |  |  |
| Q | GE2292 | t2114 | **vps-39** | V | 14035713 | G | A | SNV | nonsense | Q754* | 926 | VPS39 | (none) |
|  | GE1937 | t2189 | **vps-39** | V | 14036143 | C | T | SNV | nonsense | W626* |  |  |  |
|  | GE2056 | t2016 | **vps-39** | V | 14037839 | G | C | SNV | nonsense | Y122* |  |  |  |
| N | GE2153 | t1773 | **wapl-1** | IV | 4444464 | C | T | SNV | nonsense | W348* | 748 | WAPL | (none) |
|  | GE2305 | t1867 | **wapl-1** | IV | 4442749-4442872 | - | - | 122-bp deletion | deletion | - |  |  |  |
| p | GE2738 | t1833 | **Y54G2A.73** | IV | 3000662 | A | T | SNV | nonsense | L341* | 380 | (none) | (none) |
|  | GE2387 | t1913 | **Y54G2A.73** | IV | 3001767 | G | A | SNV | nonsense | R252* |  |  |  |
|  | GE2884 | t1755 | **Y54G2A.73** | IV | 3008481 | C | T | SNV | splice site | - |  |  |  |
| gene-23 | GE1713 | t1433 | **ZK688.9** | III | 7882477 | C | T | SNV | nonsense | W135* | 281 | TIPRL | (none) |
|  | GE2621 | t1587 | **ZK688.9** | III | 7882717 | C | T | SNV | splice site | - |  |  |  |
| gene-14 | GE2348 | t1518 | **zyg-8** | III | 12063671 | C | T | SNV | nonsense | R312* | 802 | DCLK1, DCLK2, DCLK3, DCX | Lissencephaly, X-linked, 1; Subcortical laminar heterotopia, X-linked [300067] |
|  | GE2362 | t1547 | **zyg-8** | III | 12063832 | G | A | SNV | splice site | - |  |  |  |

<sup>†</sup> Amino acid position and size derived from the longest transcript (wormbase.org, version WS275)
<sup>‡</sup> Phenotypes retrieved from omim.org

**Table 3.** Genes of interest and associated phenotypes

| Strain | Allele | Gene Name | Protein Function<sup>†</sup> | Amino Acid Change<sup>†</sup> | RNAi Phenotype<sup>‡</sup> | Mutant Phenotype | Embryonic Osmotic Integrity Defect |
| --- | --- | --- | --- | --- | --- | --- | --- |
| GE1936 | t1738 | **atg-7** | E1 ubiquitin-activating-like enzyme orthologous to the autophagic budding yeast protein Apg7p | W311* | growth variant; dauer body morphology variant; pathogen induced death increased; P granule localization defective; dauer development variant; protein aggregation variant; shortened life span; transgene subcellular localization variant; transgene expression variant; necrotic cell death variant; autophagy variant; antibody staining reduced | dead embryos | no |
| GE1958 | t1726 |  |  | Q367* |  | dead embryos | no |
| GE2627 | t1603 | **bckd-1A** | Predicted mitochondrial protein with alpha-ketoacid dehydrogenase activity | W150* | shortened life span; small | dead embryos | yes |
| GE2206 | t1514 |  |  | Q161* |  | dead embryos | yes |
| GE2840 | t1860 | **C34D4.4** | Predicted to have the following domain: Golgi apparatus membrane protein TVP23-like | W131* | -- | unfertilized oocytes | N/A |
| GE2890 | t1821 |  |  | W101* |  | unfertilized oocytes | N/A |
| GE2734 | t2029 | **C56A3.8** | Predicted to have 1-phosphatidylinositol 4-kinase activity | G62E | larval lethal; accumulated germline cell corpses; germ cell compartment morphology variant; germline nuclear positioning variant; larval arrest; cell membrane organization biogenesis variant; embryonic lethal; rachis narrow; apoptosis variant; maternal sterile; reduced brood size | unfertilized oocytes | N/A |
| GE2487 | t2149 |  |  | P82L |  | unfertilized oocytes | N/A |
| GE2886 | t2055 |  |  | E243K |  | unfertilized oocytes | N/A |
| GE2122 | t2007 | **cept-2** | Predicted to have diacylglycerol cholinephosphotransferase activity and ethanolaminephosphotransferase activity | splice site | fat content reduced; embryonic lethal; long | dead embryos | no |
| GE2047 | t2021 |  |  | W128* |  | no eggs laid (dead embryos) [ts] | some |
| GE2275 | t1517 | **cls-2** | Member of the CLASP family of microtubule-binding proteins | R102Q | locomotion variant; mitosis variant; univalent meiotic chromosomes; no polar body formation; chromosome segregation variant karyomeres early emb; mitotic chromosome segregation variant; mitotic spindle defective early emb; chromosome segregation variant; embryonic lethal; meiotic spindle defective; meiotic progression during oogenesis variant; exploded through vulva; reduced brood size; antibody subcellular localization variant; meiotic chromosome segregation variant | dead embryos | N/T |
| GE2357 | t1527 |  |  | G114R |  | dead embryos | no |
| GE1938 | t1742 | **cpt-2** | Carnitine palmitoyl transferase | W194* | embryonic lethal | dead embryos | no |
| GE2447 | t1879 |  |  | Q141* |  | dead embryos | no |
| GE2407 | t1906 | **D2096.12** | -- | L638* | locomotion variant | dead embryos | some |
| GE2499 | t1877 |  |  | Q126* |  | dead embryos | yes |
| GE2063 | t2042 | **dgtr-1** | Acyl chain transfer enzyme | G310R | sterile; sick; oocyte number decreased; germline nuclear positioning variant; oocyte septum formation variant; embryonic lethal; embryo osmotic integrity defective early emb; oocyte morphology variant; pachytene region organization variant; reduced brood size; germ cell compartment expansion variant; oogenesis variant | dead embryos | some |
| GE2135 | t2043 |  |  | splice site |  | dead embryos | yes |
| GE2541 | t2035 | **dlat-1** | Predicted to have dihydrolipoyllysine-residue acetyltransferase activity | P83L | embryonic lethal; slow growth; receptor mediated endocytosis defective; pattern of transgene expression variant; sterile progeny; transgene expression increased; general pace of development defective early emb | dead embryos | no |
| GE2335 | t2056 |  |  | Q419* |  | dead embryos | no |
| GE2402 | t1940 | **F21D5.1** | Predicted to have phosphoacetyl-glucosamine mutase activity | A436V | sterile; germ cell compartment size variant; rachis wide; rachis morphology variant; accumulated germline cell corpses; germ cell compartment morphology variant; germline nuclear positioning variant; embryonic lethal; embryo osmotic integrity defective early emb; apoptosis variant; reduced brood size; oogenesis variant | dead embryos | yes |
| GE2445 | t1935 |  |  | L539F |  | dead embryos | yes |
| GE2881 | t1744 | **F56D5.2** | -- | S107F | -- | unfertilized oocytes | N/A |
| GE2837 | t1791 |  |  | Q214* |  | unfertilized oocytes | N/A |
| GE2091 | t1772 | **nstp-2** | Predicted to have UDP-N-acetylglucosamine and UDP-xylose transmembrane transporter activity | L277H | lysosome-related organelle morphology variant; transgene subcellular localization variant; RAB-11 recycling endosome localization variant; RAB-11 recycling endosome morphology variant | dead embryos | no |
| GE2288 | t1835 |  |  | G131R |  | dead embryos | no |
| GE2391 | t1932 | **perm-5** | Predicted to have lipid binding activity | C454S | sterile; apoptosis reduced; oocytes lack nucleus; oocyte number decreased; germ cell compartment morphology variant; germline nuclear positioning variant; germ cell compartment anucleate; oocyte septum formation variant; cell membrane organization biogenesis variant; embryonic lethal; embryo osmotic integrity defective early emb; oogenesis variant; diplotene region organization variant | dead embryos | yes |
| GE2453 | t1900 |  |  | S323P |  | dead embryos | yes |
| GE2827 | t1786 | **T22B11.1** | -- | W35* | -- | unfertilized oocytes [ts] | N/A |
| GE2895 | t1866 |  |  | W356* |  | unfertilized oocytes [ts] | N/A |
| GE2399 | t1559 | **top-3** | Exhibits DNA topoisomerase type I (single strand cut, ATP-independent) activity | G59R | chromosome morphology variant; hermaphrodite germline proliferation variant; antibody staining increased; somatic gonad development variant; gonad degenerate; chromosome instability; germ cell mitosis variant; gonad arm morphology variant; meiosis variant; oocyte morphology variant; nuclear appearance variant; fewer germ cells; oogenesis variant | dead embryos | no |
| GE2220 | t1516 |  |  | Q602* |  | dead embryos | no |
| GE2512 | t1909 | **trcs-1** | Putative arylacetamide deacetylase and microsomal lipase | E373K | apoptosis reduced; diplotene absent during oogenesis; oocyte number decreased; embryo osmotic integrity defective early emb; rachis narrow; chromosome condensation variant; pachytene region organization variant; membrane trafficking variant; pachytene progression during oogenesis variant; apoptosis fails to occur; egg laying variant; germ cell compartment expansion absent; embryonic lethal; cell membrane organization biogenesis variant; no oocytes; germ cell compartment expansion variant | dead embryos [leaky ts] | yes |
| GE1939 | t1745 |  |  | Q242* |  | no eggs laid (dead embryos) [ts] | yes |
| GE2884 | t1755 | **Y54G2A.73** | -- | splice site | -- | unfertilized oocytes | N/A |
| GE2387 | t1913 |  |  | R252* |  | unfertilized oocytes | N/A |
| GE2738 | t1833 |  |  | L341* |  | unfertilized oocytes | N/A |
| GE1713 | t1433 | **ZK688.9** | Predicted to have the following domain: TIP41-like protein (TOR signaling pathway regulator) | W135* | egg laying variant; locomotion variant | dead embryos | no |
| GE2621 | t1587 |  |  | splice site |  | dead embryos | no |

[ts] = temperature sensitive
N/A = not applicable
N/T = not tested
-- = no information available
<sup>†</sup> From WormBase (WS275; wormbase.org); amino acid position derived from the longest transcript
<sup>‡</sup> Phenotypes retrieved from GExplore (genome.sfu.ca/gexplore)

### Validation of candidate gene assignments

After isolation, the mutant alleles were each localized to a chromosomal region through deficiency mapping. This data was used to corroborate the candidate gene identities derived from WGS analysis and to resolve complementation groups with more than one gene candidate. For the majority of complementation groups, the genomic position of the assigned gene was in agreement with the deficiency genetic mapping data (Figure 1).

**Figure 1.**
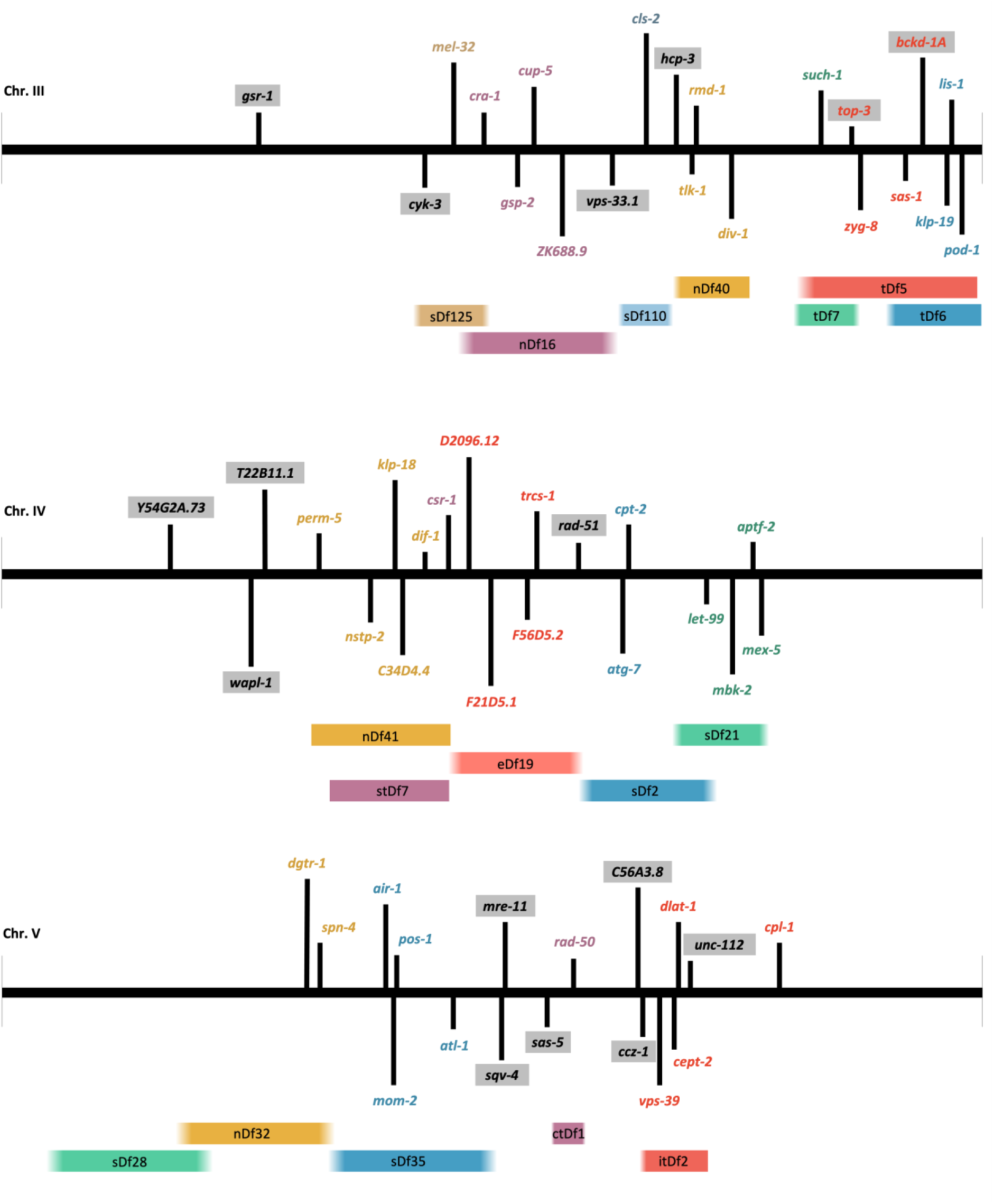
Schematic of gene assignments and deficiency mapping. Genes and deficiencies are shown with their relative positions on chromosomes III, IV, and V. Approximate boundaries of each deficiency were determined by the coordinates of the closest gene known to lie outside of the deletion, when possible (indicated by a faded edge). If no such genes with physical coordinates are known, the outermost gene known to lie inside the deletion was used as the boundary (indicated by a sharp edge). Gene names are coloured according to the deficiency under which the alleles were mapped. Genes names assigned to alleles that did not map under any of the tested deficiencies are highlighted in grey. *top-3* and *bckd-1A* on chromosome III are represented by multiple complementation groups with conflicting results from deficiency mapping.

There were some conflicts between the deficiency mapping data and the gene candidates proposed through WGS analysis. Three complementation groups that were found to not map under any of the tested deficiencies were assigned gene candidates whose genomic coordinates fall into regions covered by the tested deficiencies (alleles of *bckd-1A*, *top-3*, and *unc-112;* Figure 1). In addition, two of these groups were assigned the same gene identity as another, purportedly distinct, complementation group (Table 4). From WGS analysis, *bckd-1A* was the initial gene candidate for two different complementation groups, yet only one of these groups had been mapped to a deletion (*tDf5*) that covers the *bckd-1A* locus. Similarly, *top-3* was the assigned gene candidate for three different complementation groups, only one of which was mapped under a deficiency (*tDf5*) encompassing that gene. By performing complementation tests with select alleles (Table 4), we concluded that the two *bckd-1A* groups are not distinct, and indeed they contain mutations in the same gene. One of the groups (gene-35) originally identified as *top-3* is a double mutant which fails to complement gene-15 (*top-3*) and gene-34 (unknown gene).

**Table 4.** Complementation tests for conflicting groups

| Original Complementation Group | Strain | Allele | Preliminary Gene Candidate | Mapped Under | Complement Test Results | Final Gene Assignment |
| --- | --- | --- | --- | --- | --- | --- |
| gene-28 | GE1742 | t1461 | *bckd-1A* | None of tested deficiencies | Fails to complement: GE2206, GE2627 | *bckd-1A* |
| gene-17 | GE2627 | t1603 | *bckd-1A* | *tDf5* | Fails to complement: GE2206, GE1742 | *bckd-1A* |
|  | GE2206 | t1514 |  | *tDf5* | Fails to complement: GE2627, GE1742 |  |
| gene-15 | GE2220 | t1516 | *top-3* | *tDf5* | Fails to complement: GE2399, GE1735<br>Complements: GE2278 | *top-3* |
|  | GE2399 | t1559 |  | *tDf5* | Fails to complement: GE2220 |  |
| gene-34 | GE2278 | t1502 | *top-3* | None of tested deficiencies | Fails to complement: GE1735<br>Complements: GE2220 | unknown gene |
| gene-35 | GE1735 | t1470 | *top-3* | None of tested deficiencies | Fails to complement: GE2278, GE2220 | double mutant: *top-3* +<br>unknown gene |

N/T = not tested

Three candidate genes (*nstp-2, C34D4.4* and *F56D5.2*) were selected for additional validation by generating a deletion of the gene in a wild-type background using CRISPR-Cas9 genome editing (Norris *et al*. 2015; Au *et al*. 2019). These genes were chosen because they were expected to be of interest to the broader research community. The deletion alleles have been verified with the PCR protocol described by Au *et al*. (2019). Guide RNA sequences and deletion-flanking sequences are listed in Supplementary Table S1. Complementation testing between the newly generated CRISPR-Cas9 deletion mutants and the legacy mutant strains confirmed that the mutations are allelic, and the genes assigned to the legacy strains are correct (Supplementary Table S1)

### Human orthologs, gene ontology, and expression patterns

Of the 58 essential genes identified, 47 genes have predicted human orthologs (Table 2). Many of these genes in humans have been implicated in disease and are associated with OMIM disease phenotypes (Online Mendelian Inheritance in Man; omim.org). BLASTp searches revealed that the set of 19 GOI contains three nematode-specific genes (*F56D5.2*, *perm-5*, and *T22B11.1*) that have homologs in parasitic species, and two uncharacterized genes (*D2096.12* and *Y54G2A.73*) that do not have significant homology outside the *Caenorhabditis* genus.

To gain insight into the functions of the identified essential genes, an overrepresentation test was used to elucidate the most prominent gene ontology (GO) terms associated with them. The Biological Process terms overrepresented in the set of 58 essential genes include such terms as organelle organization (GO:0006996), nuclear division (GO:0000280), cellular metabolic process (GO:0044237), and DNA repair (GO:0006281), as shown in Figure 2. In the Molecular Function category, binding (GO:0005488) and catalytic activity (GO:0003824) are overrepresented by 41 genes (adjusted p=1.2E-07) and 28 genes (adjusted p=1.8E-03), respectively.

**Figure 2.**
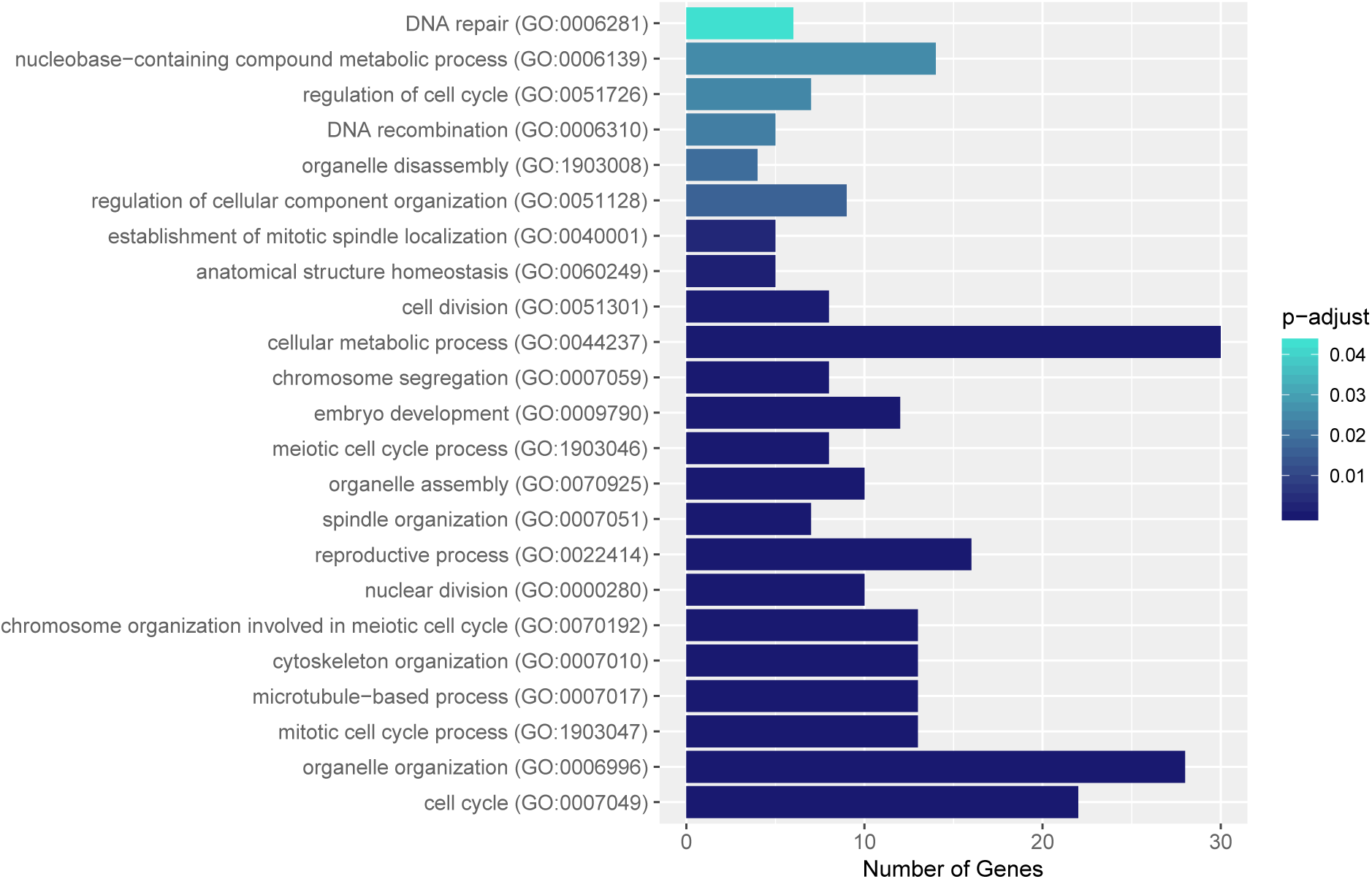
Biological Process GO terms overrepresented in the set of 58 identified essential genes. Bar length represents the number of genes in the set associated with each GO term. Overrepresentation was analyzed using PANTHER version 16.0 (Thomas *et al*. 2003) and p-values were adjusted with the Bonferroni multiple testing correction. Results were filtered to include terms with adjusted p<0.05 and edited to exclude redundant terms.

To examine the timing of gene expression throughout the life cycle, gene expression data from the modENCODE project (Hillier *et al*. 2009; Gerstein *et al*. 2010, 2014; Boeck *et al*. 2016) was retrieved from GExplore (genome.sfu.ca/gexplore; Hutter *et al*. 2009; Hutter and Suh 2016) for the 19 GOI (Supplementary Appendix S2). These data show a U-shaped expression pattern for ten of the GOI, with high expression occurring in the early embryonic stages as well as in adulthood, and particularly in the hermaphrodite gonad. This U-shaped pattern is characteristic of a maternal-effect gene, for which gene products are passed on to the embryo from the parent. Five genes have a maternal gene expression pattern as well as expression throughout other stages of the life cycle, indicating an additional, zygotic role for the gene. Seven genes have elevated expression levels in males and L4-stage hermaphrodites. These genes are suspected to be involved in sperm production or fertilization, and the associated strains were subjected to mating assays (see below).

### Temperature sensitivity and mating assays for genes of interest

The 40 alleles associated with the 19 GOI were further examined to gain insight into the phenotypic consequences of their mutations. Each allele was assayed for temperature sensitivity, as some of the original mutant screening was carried out at 25°C. Five alleles (marked with a [ts] phenotype in Table 3) were deemed temperature sensitive and could proliferate as homozygotes at a permissive temperature of 15°C, while being maternal-effect lethal or sterile at a restrictive temperature of 25°C. Curiously, four of these temperature sensitive alleles were the results of stop codons, not missense mutations.

Seven candidate genes (16 alleles) were hypothesized to be involved in male fertility, based on the production of unfertilized oocytes by hermaphrodites and/or predominantly male gene expression patterns. These 16 strains were assayed for their ability to be rescued through mating with wild-type males. 14 of the strains were rescued by the mating assay, while two strains failed to rescue (Table 5). Phenotypic rescue through mating was consistent among alleles of the same gene in five of the seven genes, while two genes had conflicting results among the pair of alleles in their complementation groups (*F56D5.2* and *nstp-2*).

**Table 5.** Putative male fertility genes

| Strain | Allele | Gene | Observed Mutant Phenotype | Successful WT Male Rescue |
| --- | --- | --- | --- | --- |
| GE2627 | t1603 | **bckd-1A** | dead embryos | yes |
| GE2206 | t1514 |  | dead embryos | yes |
| GE2840 | t1860 | **C34D4.4** | unfertilized oocytes | yes |
| GE2890 | t1821 |  | unfertilized oocytes | yes |
| GE2734 | t2029 | **C56A3.8** | unfertilized oocytes | yes |
| GE2487 | t2149 |  | unfertilized oocytes | yes |
| GE2886 | t2055 |  | unfertilized oocytes | yes |
| GE2881 | t1744 | **F56D5.2** | unfertilized oocytes | no |
| GE2837 | t1791 |  | unfertilized oocytes | yes |
| GE2091 | t1772 | **nstp-2** | dead embryos | no |
| GE2288 | t1835 |  | dead embryos | yes |
| GE2827 | t1786 | **T22B11.1** | unfertilized oocytes [ts] | yes |
| GE2895 | t1866 |  | unfertilized oocytes [ts] | yes |
| GE2884 | t1755 | **Y54G2A.73** | unfertilized oocytes | yes |
| GE2387 | t1913 |  | unfertilized oocytes | yes |
| GE2738 | t1833 |  | unfertilized oocytes | yes |

[ts] = temperature sensitive

### Terminal phenotypes of maternal-effect lethal embryos

Using DIC microscopy, the terminal phenotypes of 28 maternal-effect lethal strains (a subset of the 40 GOI strains) were observed. Representative images were selected and compiled into a catalogue of terminal phenotypes (Supplementary Appendix S3). Ten strains showed an osmotic integrity defective (OID) phenotype (as described in Sönnichsen *et al*. 2005) in nearly all embryos after incubation in distilled water, while three additional strains had only some embryos that exhibited this phenotype (Table 3). The OID phenotype was evident in embryos that filled the eggshell completely (for example, *dgtr-1*(*t2043*), Figure 3A) and eggs that burst in their hypotonic surroundings. Early embryonic arrest was observed in embryos from the two *dlat-1* mutant strains (*t2035* and *t2056*), which arrested most often with only one to four cells (for example, Figure 3B). Eleven strains had embryos that terminated with approximately 100-200 cells (for example, *ZK688.*9(*t1433*), Figure 3C); while four strains developed into two- or three-fold stage embryos that did not hatch and exhibited clear morphological defects, such as *nstp-2*(*t1835*) with a lumpy body wall and constricted nose tip (Figure 3D).

**Figure 3.**
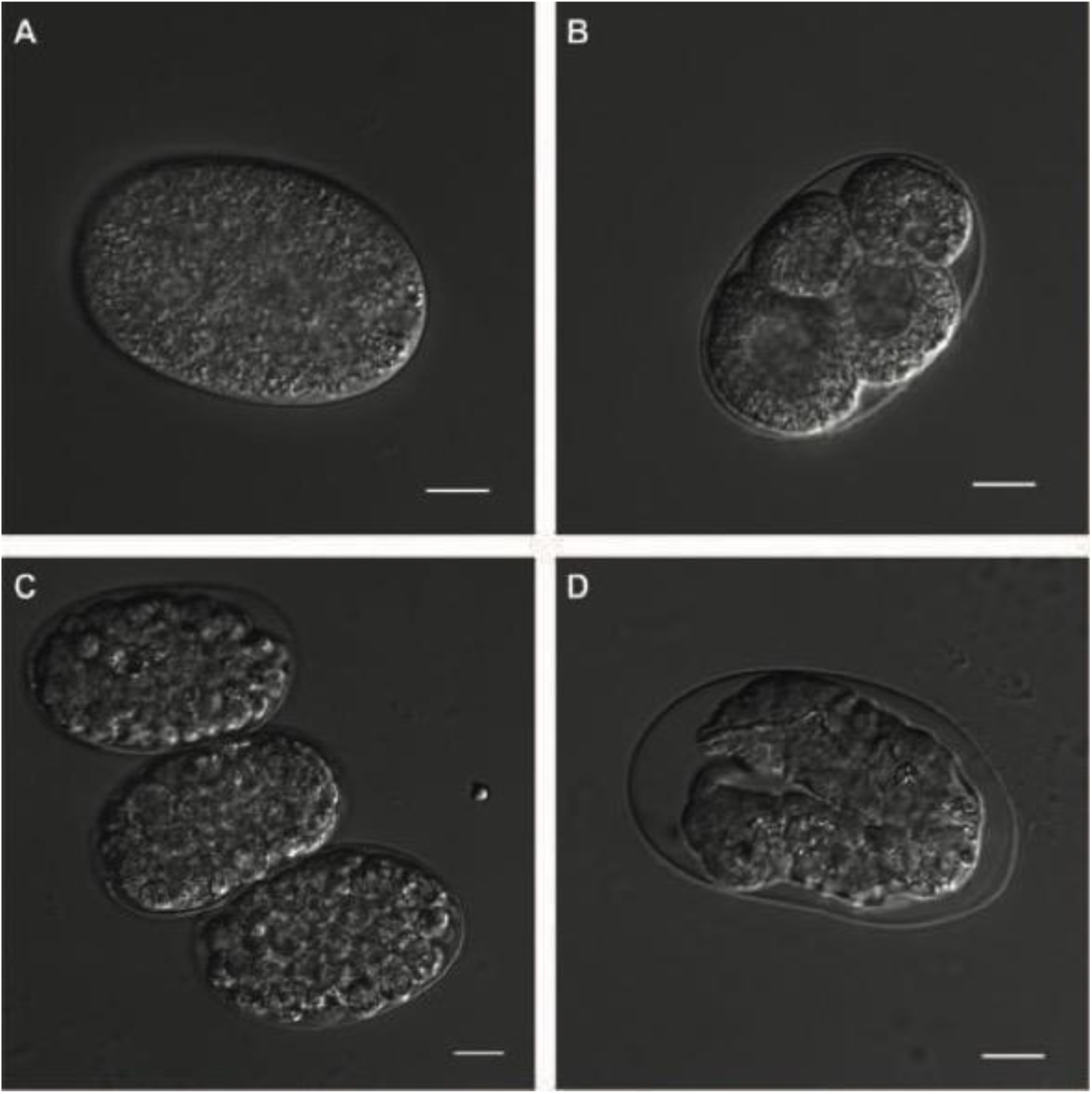
Embryonic arrest visualized with DIC microscopy for select maternal-effect lethal mutants. Eggs were dissected from homozygous mutants and imaged immediately (A) or incubated in distilled water overnight before imaging (B, C, and D). (A) Eggs dissected from *dgtr-1*(*t2043*) homozygotes exhibit signs of an osmotic integrity defect, by filling the eggshell completely. (B) *dlat-1*(*t2035*) embryos exhibit early embryonic arrest, with most embryos consisting of four cells or less. (C) *ZK688.*9(*t1433*) embryos arrest with approximately 100 cells. (D) Terminal embryos of *nstp-2*(*t1835*) have a lumpy body wall morphology and constricted nose; most animals were moving inside the eggshell but did not hatch. All scale bars represent 10 μm.

## DISCUSSION

### Revisiting legacy mutant collections with whole genome sequencing

In this study, we focused on reexamining legacy collections of *C. elegans* mutants isolated before the complete genome sequence was published (The C. elegans Sequencing Consortium 1998) and long before massively parallel sequencing was widely available. With major advances in sequencing technology in the past 30 years (reviewed in Goodwin *et al*. 2016), WGS has become affordable and accessible, making it possible to revisit past projects with new approaches and advanced capabilities. We have sequenced paired alleles from 75 complementation groups on chromosomes III, IV, and V, from which we identified 58 essential genes (Table 2).

While WGS is a powerful tool, it does not stand alone as a solution to identifying mutant alleles. This study has shown the power of having multiple alleles in a complementation group when faced with the abundance of genomic variants found in WGS analysis. Indeed, when we sequenced four single alleles, which had no complementation pairs, we were unable to designate a single mutation as the variant responsible for maternal-effect lethality (data not shown). Our approach to gene identification proved to be effective and was validated by a combination of different methods. The blind test set of 17 previously sequenced alleles from which eight of nine genes were readily identified serves as an important validation of our analysis pipeline and gives confidence in the results we obtained. In addition, the deficiency mapping data, gene expression patterns from the modENCODE project, GO term analysis, and phenotypes documented from previous experiments provide evidence to support the gene identities we assigned in these mutant collections.

The CRISPR-Cas9 deletion alleles we generated for selected gene candidates provide additional validation and will be made available to the research community to serve as useful tools for future studies. While the mutant alleles from the original study have been outcrossed, the genetic balancer background and additional mutations that persist can complicate phenotypic analysis. In contrast, these new CRISPR-Cas9 deletion strains were made in a wild-type background, which makes it much easier to handle them and interpret their mutant phenotypes. Furthermore, the pharyngeal GFP expression introduced by the gene editing approach acts as a dominant and straightforward marker for tracking the alleles in a heterozygous population. This is useful as the homozygous animals do not produce viable progeny.

The complementation groups that could not be assigned gene identities in our analysis may have been complicated by variants in noncoding regions, poor sequencing coverage, or inaccurate complementation pairing, among other possibilities. In future work, tracking down the genes we were unable to identify will require repeating complementation tests and re-tooling the analysis approach.

### Gene ontology analysis reveals common themes and gaps in our knowledge

The underlying biological themes of the 58 essential genes were revealed by examining their GO terms. The biological processes represented in Figure 2 help to confirm the nature of this set, as a collection of genes that are required for essential functions such as cell division, metabolism, and development. Performing GO-term analysis also revealed that a number of the genes in this collection lacked sufficient annotation to be interpreted this way. We found four genes about which there is little to nothing known (*D2096.12*, *F56D5.2*, *T22B11.1*, and *Y54G2A.73*). For example, *F56D5.2* is a gene with no associated GO terms, no known protein domains, and no orthologs in other model organisms. These wholly uncharacterized genes are intriguing candidates which may help uncover new biological processes and biochemical pathways that are evidently fundamental to life for this organism.

### Examining expression patterns leads to discovery of genes involved in male fertility

The life stage-specific expression patterns (Supplementary Appendix S2) provide some insight into the roles the genes in this collection play in development. 15 of the 19 GOI are highly expressed in the early embryo and hermaphrodite gonad, which suggests that the gene product is passed on to the embryo from the parent. Five of these maternal genes also have elevated expression during late embryonic and larval stages, which suggests they are pleiotropic. The zygotic functions of these genes must be non-essential or else a zygotic lethal, rather than maternal-effect lethal, phenotype would be observed.

We also identified four genes that are most highly expressed in males and L4 hermaphrodites, as well as three genes that have prominent male expression in addition to characteristic maternal expression patterns. Mating assays confirmed that these male-expressed genes have an essential role in male fertility. Studies have shown that genes expressed in sperm are largely insensitive to RNAi (Fraser *et al*. 2000; Gönczy *et al*. 2000; Reinke *et al*. 2004; del Castillo-Olivares *et al*. 2009; Zhu *et al*. 2009; Ma *et al*. 2014), making these types of genes particularly difficult to identify in high-throughput RNAi screens. With the availability of RNA-seq data across different life stages for nearly every gene in the *C. elegans* genome (Hillier *et al*. 2009; Gerstein *et al*. 2010, 2014; Boeck *et al*. 2016; Tintori *et al*. 2016; Packer *et al*. 2019), screening for characteristic gene expression patterns may be a useful approach for identifying sterile and maternal-effect lethal genes that remain to be discovered.

We propose that the seven male-expressed genes are involved in sperm production and/or function (see Table 5). These genes are mostly uncharacterized, and this is the first reporting of their involvement in male fertility. While the mutant hermaphrodites lay unfertilized oocytes (5 genes) or dead eggs (2 genes), this phenotype could be rescued in 14 of the 16 alleles by the introduction of wild-type sperm through mating. The two alleles that could not be rescued had allele pairs in the same complementation groups that were rescued in the mating assay. One of these discrepancies, between *F56D5.2*(*t1744*) and *F56D5.2*(*t1791*), was resolved when we found a second mutation in a nearby essential gene that was likely responsible for the inability of one strain to be rescued (data not shown). The presence of additional lethal mutations in the genome is unsurprising given the nature of chemical mutagenesis, and it reinforces the advantage of having multiple alleles for a gene when interpreting mutant phenotypes.

### Interpreting terminal phenotypes of maternal-effect lethal mutants

The catalogue of terminal phenotypes (Supplementary Appendix S3) created in this study provides a window into the roles the maternal-effect genes play in development. Some of these phenotypes corroborate previously observed phenotypes from RNAi studies. For example, RNAi knockdown experiments have shown that DLAT-1 is an enzyme involved in metabolic processes required for cell division in one-cell *C. elegans* embryos (Rahman *et al*. 2014). We uncovered two alleles of *dlat-1* in this study (*t2035* and *t2056*) in which most embryos arrest at the one- to four-cell stage (Figure 3B). The mutant alleles presented here can confirm previously reported phenotypes and serve as new genetic tools for continuing the study of essential gene function.

We also identified alleles for six genes that exhibit an osmotic integrity defective (OID) phenotype, resulting in embryos that filled the eggshell completely or burst in distilled water. More than 100 genes have been identified in RNAi screens as important for the osmotic integrity of developing embryos (reviewed in Stein and Golden 2018). Some of these genes have roles in lipid metabolism (Rappleye *et al*. 2003; Benenati *et al*. 2009), cellular trafficking (Rappleye *et al*. 1999), and chitin synthesis (Johnston *et al*. 2006). Four of the six genes identified with OID mutants in this study have been previously implicated in osmotic sensitivity: *dgtr-1* is involved in lipid biosynthesis (Carvalho *et al*. 2011; Olson *et al*. 2012), *trcs-1* is involved in lipid metabolism and membrane trafficking (Green *et al*. 2011); *perm-5* is predicted to have lipid binding activity; and *F21D5.1* is an ortholog of human PGM3, an enzyme involved in the hexosamine pathway which generates substrates for chitin synthase. We found OID mutants for two additional genes that were not previously characterized with this phenotype, *bckd-1A* and *D2096.12*. *bckd-1A* is a component of the branched-chain alpha-keto dehydrogenase complex, which is involved in fatty acid biosynthesis (Kniazeva *et al*. 2004); this may be indicative of a role in generating or maintaining the lipid-rich permeability barrier. *D2096.12* is a *Caenorhabditis*-specific gene with no known protein domains. Elucidating the function of this uncharacterized gene may lead to new insights about the biochemistry of eggshell formation and permeability in *C. elegans* embryos.

Most of the mutant strains we examined with DIC microscopy arrested around the 100- to 200-cell stage as a seemingly disorganized group of cells (for example, Figure 3C). Others developed into two-fold or later stage embryos that moved inside the eggshell but did not hatch (for example, Figure 3D). The terminal phenotypes documented here reveal how long the embryo can persist without the maternal contribution of gene products, and the developmental defects that ensue. Future studies might make use of fluorescent markers and automated cell lineage tracking (for example, Thomas *et al*. 1996; Schnabel *et al*. 1997; Bao *et al*. 2006; Wang *et al*. 2019) as well as single-cell transcriptome data (Tintori *et al*. 2016; Packer *et al*. 2019) to further investigate these essential genes.

### Relevance beyond *C. elegans*

In this collection of 58 essential genes, there are 47 genes (81%) with human orthologs; a two-fold enrichment when compared to all *C. elegans* genes, 41% of which have human orthologs (Kim *et al*. 2018). This is in line with previous findings that essential genes are more often phylogenetically conserved than non-essential genes (Hughes 2002; Jordan *et al*. 2002; Georgi *et al*. 2013). Essential genes in model organisms are often associated with human diseases (Culetto and Sattelle 2000; Silverman *et al*. 2009; Dickerson *et al*. 2011; Qin *et al*. 2018), making the alleles identified in this study potentially relevant to understanding human health. Indeed, there are OMIM disease phenotypes associated with a number of the human orthologs identified in Table 2. Novel mutant alleles in *C. elegans* may help us better understand genetic disorders by providing new opportunities to interrogate gene function, explore genetic interactions, and screen prospective therapeutics.

Nematode-specific genes that are essential are important to nematode biology in general and are particularly relevant in parasitic nematology. We found three genes in our GOI list (*F56D5.2*, *perm-5*, and *T22B11.1*) that have orthologs in parasitic nematode species and not in other phyla. With growing anthelminthic drug resistance around the world (Jabbar *et al*. 2006), novel management strategies are needed to combat parasitic nematodes, which infect crops, livestock, and people worldwide (Nicol *et al*. 2011; Wolstenholme *et al*. 2004; Hotez *et al*. 2008). Essential genes are desirable targets for drug development, yet identifying such genes in parasites experimentally is difficult (Kumar *et al*. 2007; Doyle *et al*. 2010). Thus, as a free-living nematode, *C. elegans* is a widely used model for genetically intractable parasitic species (Bürglin *et al*. 1998; Hashmi *et al*. 2001). Our identification of novel essential genes with orthologs in parasitic nematodes may provide new opportunities to explore management strategies.

It is our hope that the alleles and phenotypes presented here will serve as a starting point and guide future research to elucidate the specific roles these genes play in embryogenesis. All of the alleles presented in this study are available to the research community through the Caenorhabditis Genetics Center (cgc.umn.edu) and we anticipate they will serve as a valuable resource in the years to come. The wealth of material uncovered in this specific legacy collection will hopefully inspire similar explorations of other frozen mutant collections.

## ACKNOWLEDGEMENTS

The authors thank Mark L. Edgley for advice and help with strain maintenance, as well as Negin Khosravi, who replicated some of the nematode assays and conducted PCR assays with *F56D5.2*(*t1744*) to reveal an additional mutation in a nearby an essential gene. This work was supported by a CIHR Canada Graduate Scholarship-Master’s (awarded to EL) and CIHR grant PJT-148549 (awarded to DGM). This work was also supported by a grant from NSERC to DGM and an R24 NIH grant 5R240D023041 (awarded to Ann Rougvie, Paul Sternberg, Geraldine Seydoux and DGM).

